# An oomycete effector protein induces shade avoidance in *Arabidopsis* and attenuates salicylate signaling by binding to host proteins of the RADICAL-INDUCED CELL DEATH1 family

**DOI:** 10.1101/137844

**Authors:** Lennart Wirthmueller, Shuta Asai, Ghanasyam Rallapalli, Jan Sklenar, Georgina Fabro, Dae Sung Kim, Ruth Lintermann, Pinja Jaspers, Michael Wrzaczek, Jaakko Kangasjärvi, Daniel MacLean, Frank L. H. Menke, Mark J. Banfield, Jonathan D. G. Jones

## Abstract

The oomycete pathogen *Hyaloperonospora arabidopsidis* (*Hpa*) causes downy mildew disease on *Arabidopsis*. During infection, *Hpa* like other biotrophic pathogens, suppresses activation of plant innate immunity by translocating effector proteins into host cells. Some of these effectors localize to the host cell nucleus where they may manipulate transcriptional reprogramming of plant defense genes. Here we report that the nuclear-localized *Hpa* effector HaRxL106, when expressed in *Arabidopsis*, induces shade avoidance and attenuates the transcriptional response to the defense signaling molecule salicylic acid. HaRxL106 interacts with RADICAL-INDUCED CELL DEATH1 (RCD1) and loss of RCD1 function renders *Arabidopsis* resilient against HaRxL106-mediated suppression of immunity. To further characterize the molecular functions of RCD1 we solved a crystal structure of RCD1’s Poly-(ADP-ribose)-Polymerase (PARP) domain and, based on non-conservation of amino acids constituting the active site of canonical PARPs, conclude that RCD1 has no PARP activity. We report that RCD1-type proteins are phosphorylated and identified histone-modifying Mut9-like kinases (MLKs) as RCD1-interacting proteins. A *mlk1,3,4* triple mutant exhibits stronger SA-induced defense marker gene expression compared to wild-type plants. Our data suggest that HaRxL106 suppresses *Arabidopsis* innate immunity by manipulating the function(s) of RCD1 in the host cell nucleus and point towards a role of RCD1 as a transcriptional co-regulator that integrates signals from light and pathogen sensors.

## Introduction

Plants rely on their innate immune system to distinguish beneficial microbes from harmful pathogens or commensal bacteria. While plant innate immunity fends off the majority of attempted infections, specialized pathogens can subvert host defenses with effector proteins that are translocated into host cells. Many pathogen effectors interfere with cellular processes that are essential for innate immunity such as formation of cell wall appositions, secretion of antimicrobial compounds, production of reactive oxygen species (ROS) or transcriptional activation of defense genes (DebRoy et al., 2004; Nomura et al., 2006; Bozkurt et al., 2011; Anderson et al., 2012; Gangadharan et al., 2013; Asai et al., 2014). Bacterial pathogens evolved specialized secretion systems to deliver effectors into host cells (Deng et al., 2017). Likewise, the fungal rice blast pathogen *Magnaporthe oryzae* employs a specialized secretion pathway to deliver host cell-targeted effectors into a host-derived membrane-rich compartment named the biotrophic interfacial complex (Khang et al., 2010; Giraldo et al., 2013). How other filamentous plant pathogens, such as oomycetes, translocate effectors into plant cells remains poorly understood (Petre and Kamoun, 2014).

Plants respond to infection by biotrophic pathogens with elevated biosynthesis of the defense hormone salicylic acid (SA). Elevated SA levels induce fluctuations in the cellular redox status culminating in activation of the NONEXPRESSOR OF PATHOGENESIS-RELATED GENE 1 (NPR1) protein (Mou et al., 2003). Export of SA from chloroplasts to the cytoplasm leads to thioredoxin-catalyzed reduction of disulfide-linked oligomeric complexes of the NPR1 protein (Tada et al., 2008). Monomeric NPR1 translocates to the nucleus where it functions as a transcriptional co-activator and is indispensable for SA-responsiveness for most SA-induced genes (Wang et al., 2006). Some biotrophic plant pathogen effectors actively suppress SA accumulation and/or SA signaling. The maize smut fungus *Ustilago maydis* produces a host cell-targeted chorismate dismutase that may suppress SA-mediated immunity by diverting the SA-precursor chorismate into the phenylpropanoid pathway (Djamei et al., 2011). The oomycete pathogen *Hyaloperonospora arabidopsidis* (*Hpa*) suppresses transcriptional upregulation of the SA marker gene *PATHOGENESIS-RELATED GENE 1* (*PR1*) in infected cells of its host *Arabidopsis thaliana* (Caillaud et al., 2013). At least two *Hpa* effector proteins interfere with SA signaling when expressed as transgenes in *Arabidopsis*. Effector HaRxL44 appears to attenuate SA signal transduction by targeting the MEDIATOR subunit Med19 for proteasomal degradation (Caillaud et al., 2013), while effector HaRxL62 interferes with SA signaling by an unknown mechanism (Asai et al., 2014). Whether pathogen effectors manipulate proteins of the NPR class directly remains unknown. However, pathogens interfere with other processes that indirectly promote SA signaling. The *Xanthomonas campestris* effector protein XopJ is a Cys protease that cleaves the RPT6 subunit of the 19S regulatory particle of the proteasome, thereby interfering with targeted degradation of poly-ubiquitinated proteasome substrates. Via this mechanism, XopJ also attenuates proteasomal turnover of NPR1, which is required for full transcriptional activation of NPR1 target genes (Üstün and Börnke, 2015; Spoel et al., 2009).

Shade avoidance in *Arabidopsis*, activated by a low red/far-red light ratio, attenuates the transcriptional response to SA (Genoud et al., 2002; de Wit et al., 2013; Gangappa et al., 2016). Notably, simulated shade conditions also suppress transcript changes induced by exogenous application of Methyl-Jasmonate (MeJA) and attenuate plant defense towards necrotrophic pathogens and herbivores (Izaguirre et al., 2006; Cerrudo et al., 2012; de Wit et al., 2013). This is remarkable given that SA- and JA-responsive gene networks are antagonistically regulated in response to infection by biotrophic and necrotrophic pathogens (Pieterse et al., 2012; Caarls et al., 2015). As a consequence, plants become more susceptible to infection by both biotrophic and necrotrophic pathogens when shade avoidance is activated either environmentally, or genetically by mutations in *PHYTOCHROME B* (*PHYB*) (Genoud et al., 2002; Izaguirre et al., 2006; de Wit et al., 2013).

*Arabidopsis* RADICAL-INDUCED CELL DEATH1 (RCD1) has also been proposed to act as a positive regulator of SA signaling. Loss of *RCD1* function does not alter SA levels but transcript levels of many NPR1 target genes are lower in *rcd1* mutants when compared to wild-type plants (Ahlfors et al., 2004; Brosché et al., 2014). RCD1 is the founding member of a plant-specific protein family characterized by a central domain with sequence similarity to the catalytic domain of Poly-(ADP-ribose)-polymerases (PARPs) (Lamb et al., 2012). In contrast to canonical PARPs that covalently modify target proteins by ADP-ribosylation, *Arabidopsis* RCD1 does not show PARP activity *in vitro* when expressed as a GST fusion (Jaspers et al., 2010a). In addition to the central PARP domain, RCD1 and its paralog SIMILAR TO RCD ONE 1 (SRO1) have an N-terminal WWE domain and a C-terminal RST domain. Proteins with this domain architecture are conserved in all land plants and are referred to as type I proteins. In contrast, the presence of additional family members that lack the N-terminal WWE domain (type II) appears to be specific to the *Brassicaceae* (Jaspers et al., 2010a). Type I proteins from *Arabidopsis* and rice localize to the plant cell nucleus and bind to many sequence-unrelated transcription factors via their RST domains (Katiyar-Agarwal et al., 2006; Jaspers et al., 2009; You et al., 2014). Therefore, RCD1 might influence SA signal transduction by interacting with transcription factors that mediate SA-induced transcriptome changes. Notably, an RCD1 homologue from wheat (TaSRO1) shows PARP activity when expressed in *E. coli*, suggesting that some RCD1-type proteins may be enzymatically active (Liu et al., 2014).

RCD1 interacts with the *Hpa* effector HaRxL106 in a yeast-two-hybrid (Y2H) assay and this effector renders *Arabidopsis* more susceptible to biotrophic pathogens when expressed as a transgene (Fabro et al., 2011; Mukhtar et al., 2011). In plant cells, HaRxL106 binds to nuclear transport receptors of the importin-α class with affinity in the low micro-molar range and is actively transported into the nucleus, indicative of a virulence-promoting activity of the effector in the host cell nucleus (Wirthmueller et al., 2015). Here we report that HaRxL106, when expressed as a transgene, affects both SA signaling and light-dependent developmental processes. We identify RCD1 as a likely virulence target of HaRxL106 and report that RCD1 interacts with histone-modifying kinases that impinge on SA signaling.

## Results

### HaRxL106-expressing *Arabidopsis* plants exhibit attenuated light and defense signaling

To characterize HaRxL106-interacting proteins from *Arabidopsis* we generated transgenic lines expressing HaRxL106 with an N-terminal YFP or 3xHA-StrepII (HS) epitope tag under control of the *35S* promoter. As previously reported for transgenic plants expressing untagged HaRxL106 (Fabro et al., 2011), these lines are hyper-susceptible to infection by the compatible *Hpa* isolate Noco2 (Fig. 1A; one-way ANOVA, F_8,171_=3.14, p=5.73x10^-16^; Tukey-Kramer post-hoc test, p<0.05). While analyzing lines expressing HaRxL106 we noticed that they show signs of constitutive shade avoidance, specifically longer hypocotyls and elongated petioles under white light conditions (Fig. 1B). Differences in hypocotyl length between wild type plants and transgenic lines were more pronounced when we grew seedlings under a lower fluence rate of white light (12 μmol m^-2^ s^-1^) (Fig. 1, C and D; one-way ANOVA, F_5,174_=215.48, p=1.42x10^-72^; Tukey-Kramer post-hoc test, p<0.05). Under these conditions HaRxL106-expressing seedlings were indistinguishable from the *phyB-9* mutant that shows constitutive shade avoidance (Reed et al., 1993). Lines expressing control constructs YFP and HS did not differ from wild type plants in hypocotyl length (Fig. 1, C and D). In contrast, differences in hypocotyl length between HaRxL106-expressing transgenic lines and wild type plants were much smaller when we grew seedlings in darkness (Fig. 1D; one-way ANOVA, F_5,174_=3.14, p=9.67x10^-3^; Tukey-Kramer post-hoc test, p<0.05). This suggests that in addition to defense signaling, either light perception or light signal transduction is attenuated in lines expressing HaRxL106.

**Figure 1.**
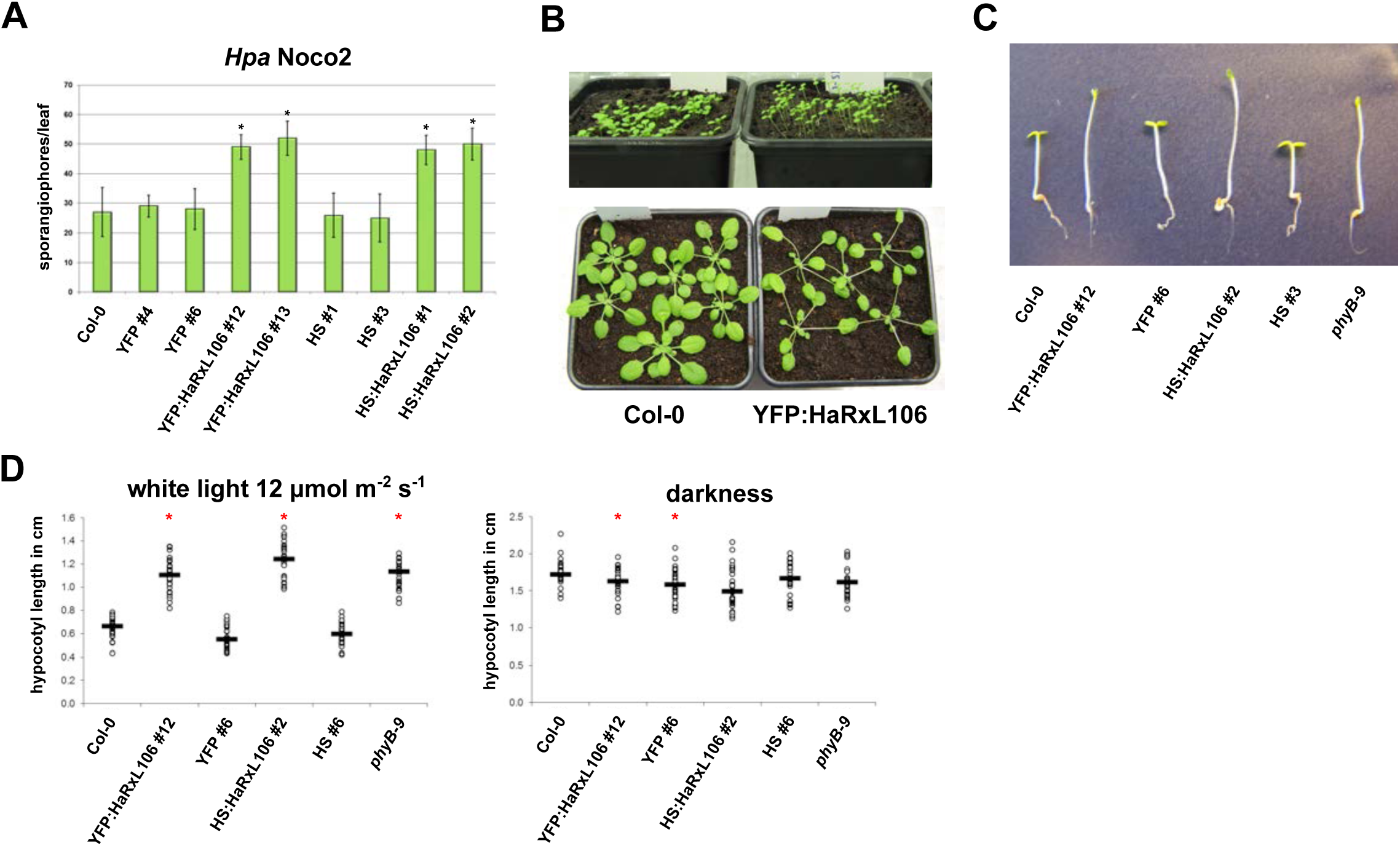
**(A)** Sporulation of the virulent *Hpa* isolate Noco2 on 6-week-old plants of the indicated genotypes quantified by the number of sporangiophores per leaf. The results shown are representative of two independent biological experiments, n=20, error bars show SE, asterisks indicate differences from Col-0 (one-way ANOVA; Tukey-Kramer post-hoc test, p<0.05). **(B)** Constitutive expression of HaRxL106 induces shade avoidance in *Arabidopsis*. The top panel shows 10-day-old seedlings of Col-0 and a representative *35S*_*Pro*_*:YFP:HaRxL106* line. Bottom panel shows 4-week-old plants from both genotypes grown under short day condition and a fluence rate of ∼120 µmol m^-2^ s^-1^. **(C)** Five-day-old seedlings of the indicated genotypes germinated under a lower fluence rate of ∼12 µmol m^-2^ s^-1^. **(D)** Quantification of seedling hypocotyl length of the indicated genotypes grown as in (C) or in darkness. The results shown are representative of three independent biological experiments, n=30, horizontal bars denote median, asterisks indicate mean values different from Col-0 (one-way ANOVA; Tukey-Kramer post-hoc test, p<0.05).

### Effector HaRxL106 suppresses SA signal transduction but not SA levels

Plants that undergo shade avoidance, either induced by supplementary FR light or by mutations in *PHYB*, show an attenuated transcriptional response to SA and MeJA (Genoud et al., 2002; de Wit et al., 2013). As suppression of SA signal transduction would be a conceivable virulence mechanism for an effector of a biotrophic pathogen, we tested SA-induced up-regulation of the SA marker gene *PR1* by qPCR in Col-0 plants and two transgenic lines expressing YFP:HaRxL106 and HS:HaRxL106, respectively (Fig. 2A). As expected, SA treatment induced *PR1* mRNA levels in Col-0 plants but not in the *npr1-1* mutant (Cao et al., 1994) (one-way ANOVA, F_7,16_=78.60, p=3.39x10^-11^; Tukey-Kramer post-hoc test, p<0.05). In contrast, *PR1* expression levels in SA-treated HaRxL106 transgenic lines were comparable to those in mock treated Col-0 plants suggesting that HaRxL106 affects either endogenous SA levels or SA signal transduction (Fig. 2A). To distinguish between these two possibilities we quantified levels of SA in wild type plants, the *sid2-1* mutant that is impaired in pathogen-triggered SA biosynthesis (Wildermuth et al., 2001), and HaRxL106 transgenics. There was a trend for lower SA levels in the *sid2-1* mutant compared to Col-0 in plants infiltrated with *Pseudomonas syringae pv. tomato* (*Pst*) DC3000 (1x10^8^ cfu/ml) 24h earlier (Fig. 2B). In contrast, SA levels in HaRxL106-expressing lines were comparable to Col-0 (experiments A and B) or intermediate between Col-0 and *sid2-1* (experiment C). When we analyzed the data using a linear mixed effects model we found no statistical differences between the genotypes (p<0.05). Despite some variability between SA measurements in the three independent biological experiments, these results suggest that HaRxL106 does not substantially alter SA levels but nevertheless strongly attenuates SA-induced transcriptional regulation of the SA marker gene *PR1*.

**Figure 2.**
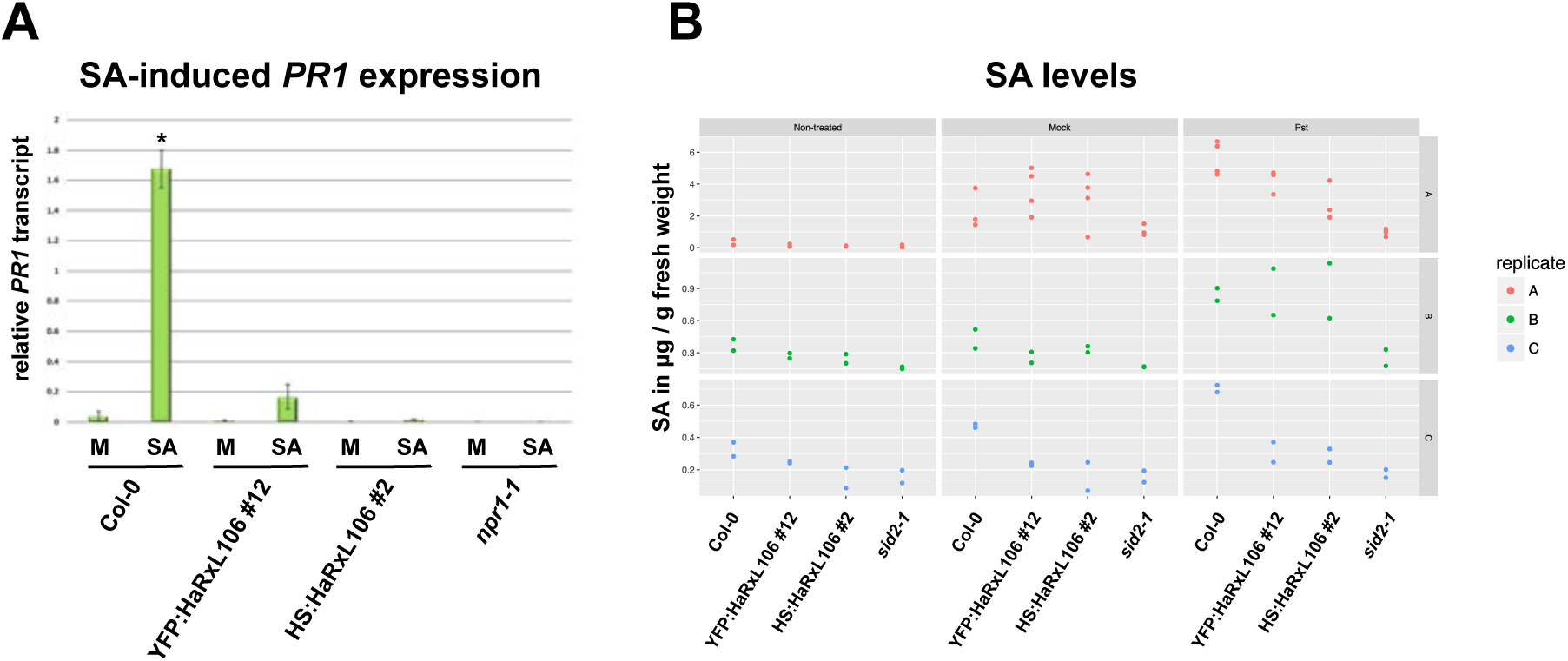
**(A)** HaRxL106 suppresses SA-induced *PR1* expression. Four-week-old plants of the indicated genotypes were sprayed with 0.1 mM SA or a mock solution and *PR1* expression levels were analyzed by qRT-PCR 8 h later. *PR1* expression levels were normalized by *EF1α* expression. The plot shows the mean of *PR1/EF1α* expression from three independent biological experiments. Error bars show SE, asterisk indicates mean value different from Col-0 mock treatment (one-way ANOVA; Tukey-Kramer post-hoc test, p<0.05). **(B)** Quantification of SA levels in the indicated genotypes under non-treated conditions and 24 h after infiltration with 10^8^ cfu/ml of *Pst* DC3000 or a 10 mM MgCl_2_ mock solution. Red, green and blue represent data from three independent biological experiments. Dots of the same color represent technical replicates.

### Effector HaRxL106 attenuates NPR1-dependent defense activation

HaRxL106 has a nuclear localization sequence (NLS) and is actively transported into nuclei of plant cells via karyopherins of the importin-α group (Wirthmueller et al., 2015). Given that NPR1 is an important nuclear signal integrator of the SA pathway, we tested whether HaRxL106 affects NPR1 localization or NPR1 protein levels. The *npr1-1* mutant has been complemented by a *35S*_*Pro*_*:NPR1:GFP* transgene under long day conditions (Kinkema et al., 2000). When the *35S*_*Pro*_*:NPR1:GFP* line is grown under short day conditions the plants show signs of constitutive defense activation including severe stunting, development of micro lesions and high expression levels of *PR1* (one-way ANOVA, F_3,8_=5.94, p=0.02; Tukey-Kramer post-hoc test, p<0.05; Fig. 3A and B; Love et al., 2012). We transformed the *35S*_*Pro*_*:NPR1:GFP* line with the *35S*_*Pro*_*:HS:HaRxL106* construct and grew independent T1 transformed lines under short day conditions. Expression of HaRxL106 completely suppressed the stunting of the *35S*_*Pro*_*:NPR1:GFP* line in 12 out of 14 transgenics (Fig. 3A). HaRxL106 also reverted the constitutive *PR1* expression of the *35S*_*Pro*_*:NPR1:GFP* line (Fig. 3B). This suppression was not due to lower NPR1:GFP protein levels as shown by the Western blot in Fig. 3C. Consistent with constitutively activated defense, we observed nuclear localization of NPR1:GFP in guard cells of plants grown under short day condition even without exogenous SA application (Fig. 3D). NPR1:GFP also localized to nuclei in double transgenic lines co-expressing HS:HaRxL106 (Fig. 3D). Taken together these results show that HaRxL106 does not attenuate SA signal transduction by altering protein levels or localization of NPR1. As HaRxL106 suppresses constitutive *PR1* gene expression induced by the *35S*_*Pro*_*:NPR1:GFP* transgene, the effector must either act on a step downstream of nuclear NPR1 signaling or disrupt the nuclear transactivator function of NPR1 itself.

**Figure 3.**
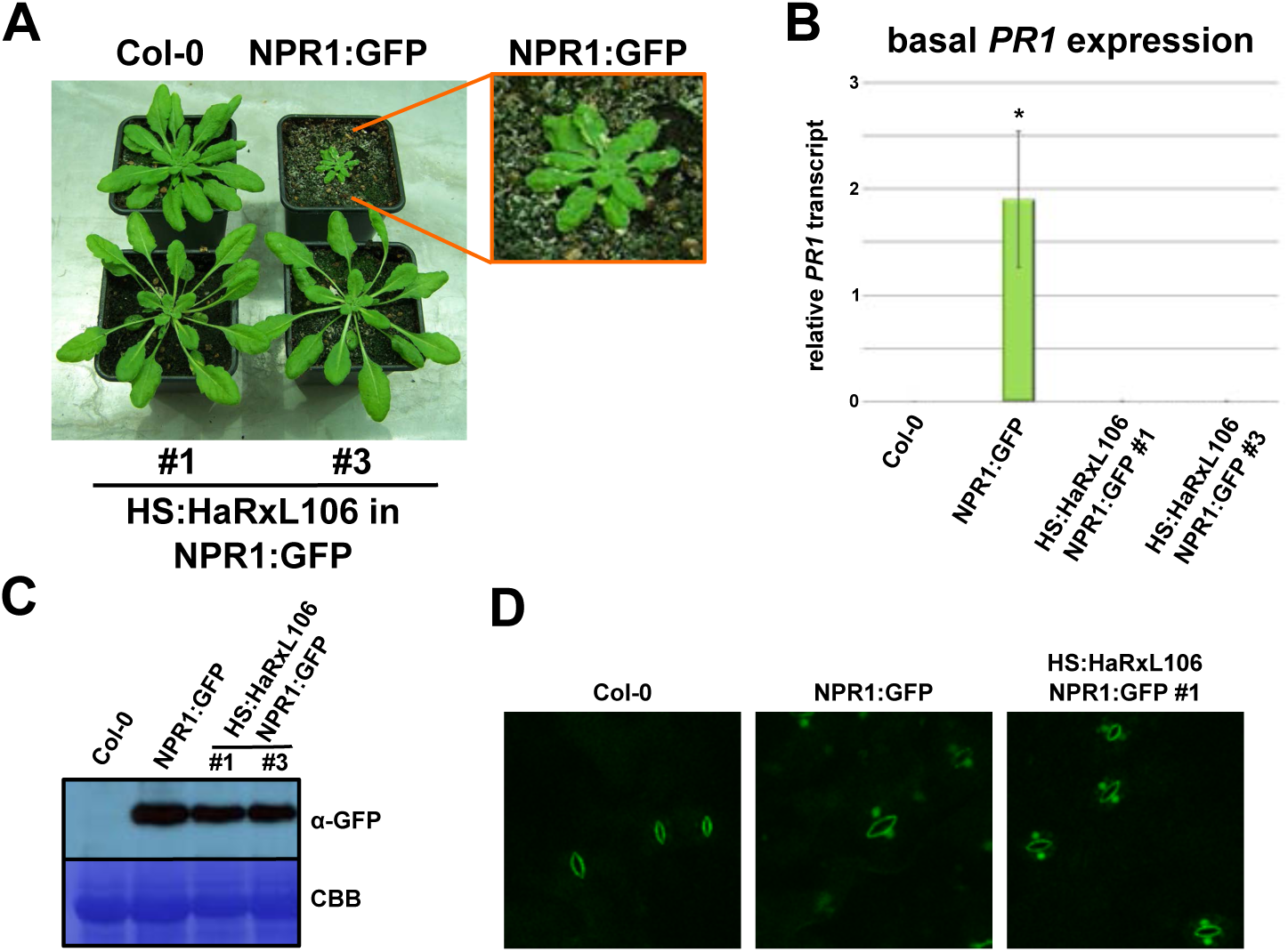
HaRxL106 suppresses constitutive defense signaling induced by NPR1:GFP over-expression under short day conditions. **(A)** Morphology of 5-week-old Col-0 and *35S*_*Pro*_*:NPR1:GFP* plants (top row) and two independent double transgenic *35S*_*Pro*_*:NPR1:GFP* lines co-expressing *35S*_*Pro*_*:HS:HaRxL106* (bottom row). The inset shows spontaneous lesions forming in *35S*_*Pro*_*:NPR1:GFP* plants. **(B)** Basal *PR1* expression in the lines shown in (A) as determined by qRT-PCR. *PR1* expression levels were normalized by *EF1α* expression. The plot shows the mean of *PR1/EF1α* expression from three independent biological experiments. Error bars show SE, asterisk indicates mean value different from Col-0 (one-way ANOVA; Tukey-Kramer post-hoc test, p<0.05). **(C)** Western blot showing accumulation of NPR1:GFP protein in the lines shown in (A). The Western blot is representative of three independent biological experiments. **(D)** Representative (n>10) confocal microscopy images showing nuclear accumulation of NPR1:GFP protein in short day conditions in *35S*_*Pro*_*:NPR1:GFP* plants and plants co-expressing *35S*_*Pro*_*:HS:HaRxL106.* The signal in Col-0 is auto-fluorescence from stomata.

### HaRxL106 over-expressing lines show a partial transcription profile overlap with the *radical-induced cell death1-1* mutant

Several *Arabidopsis* proteins that interact with HaRxL106 in the yeast-two-hybrid (Y2H) system have been reported (Mukhtar et al., 2011). HaRxL106 interactors identified included several importin-α-type karyopherins, the tri-helix transcription factor ARABIDOPSIS 6b-INTERACTING PROTEIN 1-LIKE1 (ASIL1), transcription factor TEOSINTE BRANCHED CYCLOIDEA AND PCF 14 (TCP14) and RCD1. We reasoned that if one or several of these proteins constitute virulence targets of HaRxL106, the transcriptome profile of the corresponding mutants might show similarities to the transcriptome profile of HaRxL106 over-expressors. The transcriptome of *tcp14* knock-out mutants has recently been analyzed and revealed a set of 18 genes that are differentially expressed in two independent *tcp14* T-DNA lines (Yang et al., 2017). We performed transcriptome profiling using the EXPRSS RNAseq pipeline (Rallapalli et al., 2014; Sohn et al., 2014) to compare the transcriptome profiles of Col-0, the HS:HaRxL106 over-expressor line #2, *rcd1-1*, *asil1-1* and the importin-α mutant *mos6-1* (Overmyer et al., 2000; Gao et al., 2009; Palma et al., 2005). To compare the transcriptional response of all lines to a biotrophic pathogen, we performed transcriptome profiling of non-treated plants as well as plants infiltrated with 5 × 10^5^ cfu/ml of *Pst* DC3000. Plants infiltrated with 10 mM MgCl_2_ served as mock control and we harvested all samples 24 h after infiltration. We applied a false discovery rate (FDR) of less than 0.001 and a two-fold change in expression as criteria to identify differentially expressed genes (DEGs) (Supplementary Table S1). As shown in Fig. 4A (and supplementary Fig. S1) the HaRxL106-expressing line showed the largest number of DEGs in comparison to Col-0 wild type with 1040, 844 and 1352 DEGs in non-treated, mock-infiltrated and *Pst*-infiltrated plants, respectively. In *rcd1-1,* between 181 and 429 genes differed in expression compared to Col-0. In contrast, analysis of the *asil1-1* and *mos6-1* mutants, as well as the published *tcp14* dataset, revealed a comparatively small number of DEGs (<48).

**Figure 4.**
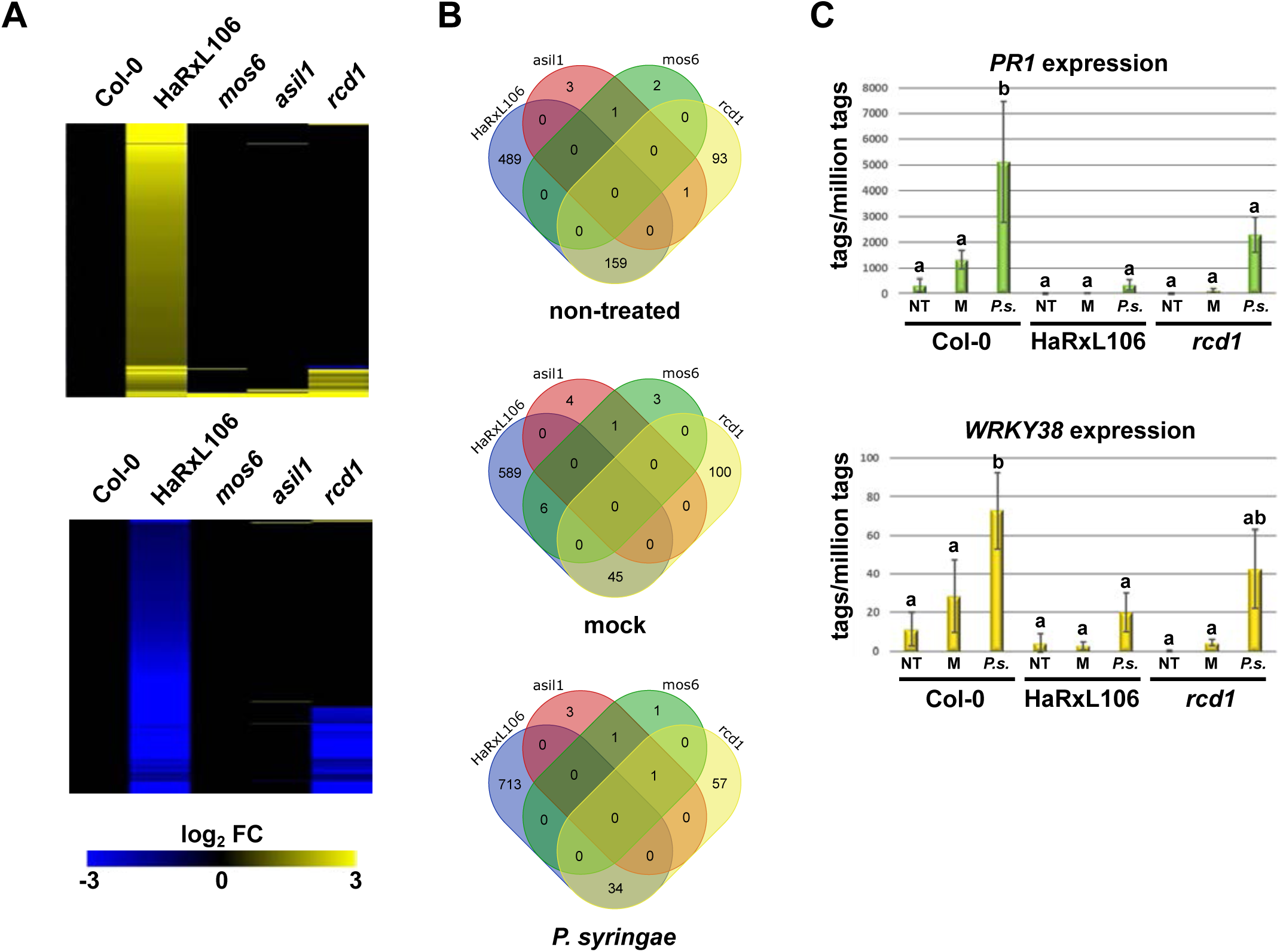
Comparative transcriptome profiling of Col-0, *35S*_*Pro*_*:HS:HaRxL106* line #2 and mutants of HaRxL106-interacting *Arabidopsis* proteins. **(A)** Comparison of transcriptome changes induced by HaRxL106 overexpression and loss of function alleles of *mos6, asil1,* and *rcd1* under non-treated conditions. Up-regulated genes in yellow, repressed genes in blue, FC = fold change. Only those genes that differ in transcript levels from Col-0 in at least one of the other genotypes are shown. **(B)** Overlap of genes repressed in HS:HaRxL106 compared to Col-0 and the transcriptome profiles of *mos6*, *asil1* and *rcd1*. For genes over-expressed in HS::HaRxL106 and all differentially expressed genes see Fig. S1. Venn diagrams were prepared using http://bioinformatics.psb.ugent.be/webtools/Venn/. **(C)** Expression profiles of the SA-marker genes *PR1* and *WRKY38* in Col-0, HS:HaRxL106 line #2 and the *rcd1-1* mutant extracted from the RNA-seq data set. The plots show mean expression values of three independent biological replicates, error bars denote SD, letters indicate differences between genotypes/treatments (one way ANOVA; Tukey-Kramer post-hoc test, p<0.05).

We noticed a partial overlap in DEGs between HaRxL106 and *rcd1-1*, particularly in the repressed genes (Fig. 4 A, B, see Fig. S1 for induced genes). In non-treated plants, 63% of genes repressed in *rcd1-1* also exhibited lower transcript abundance in the HaRxL106 transgenic line. In mock-treated and *Pst*-infiltrated tissue this overlap was substantially lower, with 31% and 37% of shared repressed genes, respectively (Fig. 4B). Vice versa, out of all genes repressed in the HaRxL106 transgenic line only 25%, 7% and 5% were also expressed at lower levels in *rcd1-1* in non-treated, mock-treated and *Pst*-infiltrated tissue, respectively (Fig. 4B). Among the genes repressed in non-treated *rcd1-1* and the HaRxL106-expressing line there was an over-representation of defense-related transcripts as determined by analysis of Gene Ontology (GO) terms (Vandepoele et al., 2009) and cis-regulatory promoter elements (O’Connor et al., 2005) (Table S1). Lower transcript abundance of SA defense genes in *rcd1-1* is consistent with a previously reported transcriptome profiling of *rcd1* mutants without pathogen challenge (Brosché et al., 2014). Figure 4C shows expression profiles of two selected SA marker genes, *PR1* and *WRKY38*. Transcript levels of both genes were induced by *Pst* infection and to lesser extent by MgCl_2_ infiltration in wild type plants. In contrast, in the HaRxL106-expressing line and in *rcd1-1*, transcriptional up-regulation of both SA marker genes by *Pst* infection or MgCl_2_ infiltration was attenuated (Fig. 4C; *PR1* one-way ANOVA, F_8,18_=5.94, p=9.5 x 10^-5^; *WRKY38* one-way ANOVA, F_8,18_=7.64, p=1.81 x 10^-4^; Tukey-Kramer post-hoc tests, p<0.05). Therefore, both loss of *RCD1* function and ectopic expression of HaRxL106, lead to repression of SA defense genes in naïve and challenged plants. The partial overlap between genes that were repressed in the HS:HaRxL106 line and *rcd1-1* prompted us to characterize the interaction between the two proteins in more detail.

### HaRxL106 interacts with RCD1 and SRO1 proteins and RCD1 quantitatively contributes to SA signal transduction

To test for interaction between HaRxL106 and RCD1 in *Arabidopsis*, we made use of a transgenic line in which the *rcd1-3* mutation is complemented by expression of an *RCD1*_*Pro*_*:RCD1:HA* construct (Jaspers et al., 2009). We transformed the RCD1:HA line with YFP:HaRxL106 and selected double transgenic lines. When we immunoprecipitated YFP:HaRxL106 from these lines, RCD1:HA co-purified with HaRxL106 whilst a cross-reacting band detected by the α-HA antibody did not (Fig. 5A). Therefore YFP:HaRxL106 interacts with functional epitope-tagged RCD1:HA protein in *Arabidopsis*.

**Figure 5.**
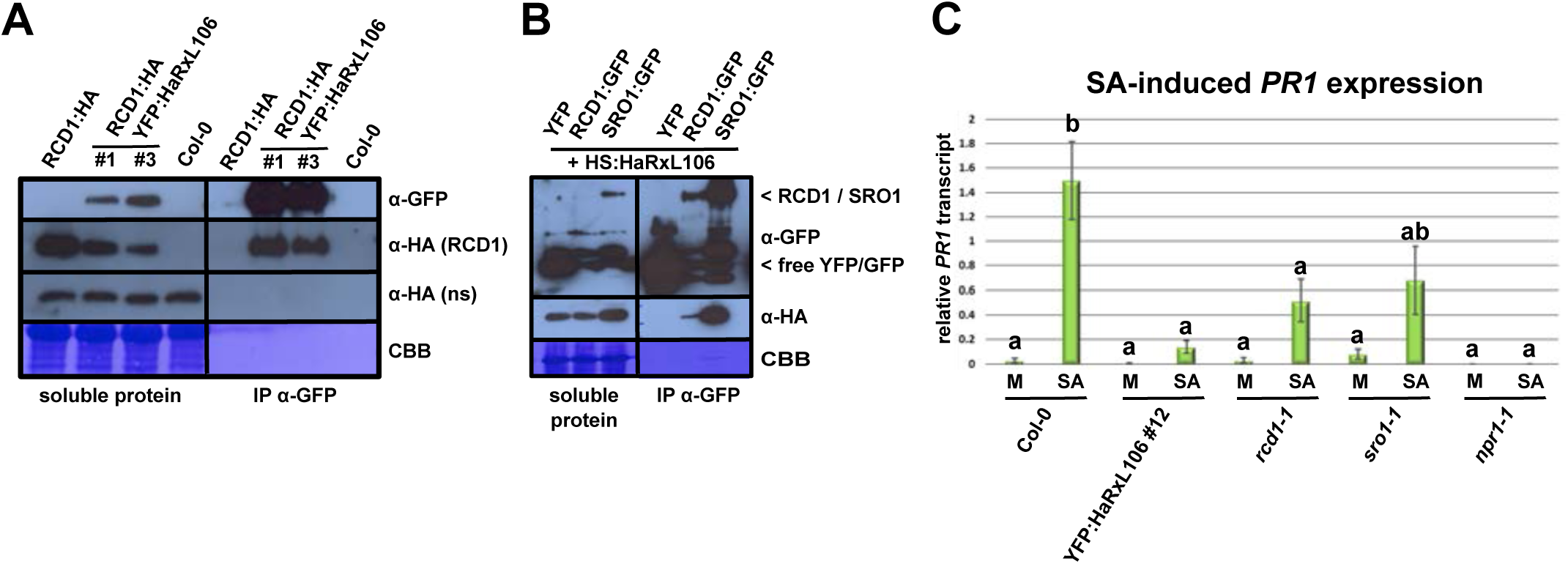
HaRxL106 interacts with RCD1 and SRO1 in plant cells and both proteins contribute to SA-induced *PR1* expression. **(A)** Functional RCD1:HA protein co-immunoprecipitates with YFP:HaRxL106 in *Arabidopsis.* YFP:HaRxL106 was immuno-precipitated from double transgenic lines expressing RCD1:HA, proteins were resolved by SDS-PAGE, transferred onto PVDF membrane and probed with the indicated antibodies. ns = non-specific band detected by the α-HA antibody. CBB = Coomassie brilliant blue stain. This result is representative of three independent biological experiments. **(B)** HS:HaRxL106 co-immunoprecipitates with GFP-tagged variants of RCD1 and SRO1 following transient expression in *N. benthamiana.* GFP-tagged proteins, or YFP as a control, were immunoprecipitated, proteins were resolved by SDS-PAGE, transferred onto PVDF membrane and probed with the indicated antibodies. Co-immunoprecipitation of HaRxL106 with RCD1 and SRO1 is based on three and two independent biological experiments, respectively. **(C)** SA-induced *PR1* gene expression in Col-0, YFP:HaRxL106 line #12, *rcd1-1*, *sro1-1* and *npr1-1* mutants. Four-week-old plants were sprayed with 0.1 mM SA or a mock solution and *PR1* expression levels were analyzed by qRT-PCR 8 h later. *PR1* expression levels were normalized by *EF1α* expression. The plot shows the mean of *PR1/EF1α* expression from five independent biological experiments. Error bars show SE, letters indicate differences between genotypes/treatments (one-way ANOVA; Tukey-Kramer post-hoc test, p<0.05).

*RCD1* and its paralog *SRO1* show unequal genetic redundancy with respect to plant development and responses to abiotic stress with *RCD1* making a stronger contribution (Jaspers et al., 2009; Teotia and Lamb, 2009). RCD1 and SRO1 share the same domain structure comprising an N-terminal WWE domain, a central PARP domain and a C-terminal RST domain (Jaspers et al., 2010b). To test whether HaRxL106 also interacts with SRO1 we used *Agrobacterium tumefaciens*-mediated transient protein expression in *Nicotiana benthamiana.* We co-expressed HS:HaRxL106 and C-terminally GFP-tagged versions of RCD1 and SRO1. Although all constructs were expressed from the strong *35S* promoter we did not detect RCD1:GFP by Western blot in total protein extracts (Fig. 5B). In contrast, SRO1:GFP and HS:HaRxL106 were detectable with the respective antibodies. As shown in Fig. 5B HS:HaRxL106 co-immunoprecipitated with both RCD1:GFP and SRO1:GFP but not with free YFP that we used as control. This suggests that HaRxL106 interacts with both RCD1 and SRO1 in plant cells.

The partial redundancy between *RCD1* and *SRO1* prompted us to test if *SRO1* also contributes to transcriptional regulation of NPR1 target genes. We compared SA-induced transcriptional upregulation of *PR1* in wild type plants, the YFP:HaRxL106 line, *rcd1-1* and *sro1-1*. *PR1* levels 8h after SA spraying were approximately 3-fold lower in *rcd1-1* and *sro1-1* but the difference was only statistically significant in the case of *rcd1-1* (one-way ANOVA, F_9,36_=7.7, p=3.32 × 10^-6^; Tukey-Kramer post-hoc test, p<0.05). The *rcd1-1* and *sro1-1* mutations had a weaker effect on *PR1* expression than the YFP:HaRxL106 transgene (Fig. 5C). Nevertheless, our data suggest that *RCD1* is required for full SA-induced *PR1* expression.

### The C-terminal 58 amino acids of HaRxL106 are required for RCD1-binding and attenuation of light and defense signaling

To test whether HaRxL106 binding to RCD1 correlates with its defense-suppressing activities we generated a mutant variant of HaRxL106 that does not bind to RCD1. The protein sequence C-terminal to the HaRxL106’s signal peptide and RxLR motif can be divided into two regions based on predicted protein secondary structure: a larger domain with a predicted α-helical WY-fold (Win et al., 2012) and a 58 amino acid C-terminal region that mediates binding to importin-α (Wirthmueller et al., 2015). To narrow down the RCD1-binding site in HaRxL106, we used the Y2H system to compare RCD1 binding to full-length HaRxL106, an HaRxL106 variant lacking the 56 C-terminal amino acids (HaRxL106 ΔC) and to the C-terminal 58 amino acids alone (HaRxL106-Cterm58). Unexpectedly, we did not detect interaction of the two full-length proteins in yeast suggesting that the sensitivity of the Y2H reporters under our conditions was lower than in Mukhtar et al., 2011. However, we found the HaRxL106 C-terminus interacts with RCD1 (Fig. 6A). Next, we tested which domain(s) of RCD1 are required for binding to the HaRxL106 C-terminus. The WWE, PARP and RST domains of RCD1 are separated by regions of 50-100 amino acids predicted to be disordered (Kragelund et al., 2012). As shown in Fig. 6A the HaRxL106 C-terminus did not interact with the isolated WWE, PARP or RST domains, nor did we detect binding to a construct encompassing PARP and RST domain. However, a fragment spanning the WWE and PARP domains activated the *HIS3* reporter when co-expressed with the HaRxL106 C-terminus. Notably, this region of RCD1 also showed a weaker interaction with full-length HaRxL106 protein (Fig. 6A). This suggests that HaRxL106 specifically binds to the RCD1 WWE-PARP domains via its C-terminal 58 amino acid peptide, and that RCD1’s RST domain might negatively affect binding of the effector to the N-terminal domains of RCD1. We next tested whether the HaRxL106 C-terminal 58 amino acids are necessary for altered light and SA signaling in *Arabidopsis*. To this end we transformed an *RFP:NLS:HaRxL106 ΔC* construct lacking the C-terminus of the effector into Col-0. Because the *HaRxL106 ΔC* construct also lacks the effector’s NLS this fusion protein carries a SV40 NLS to ensure efficient nuclear import (Wirthmueller et al., 2015). As controls we also generated transgenic RFP:HaRxL106 lines and lines expressing a fusion of RFP and the 58 C-terminal amino acids of HaRxL106 (RFP:HaRxL106-Cterm58). All constructs were under control of the *35S* promoter and we confirmed expression of the RFP fusion proteins by Western blot (Fig. 6B).

**Figure 6.**
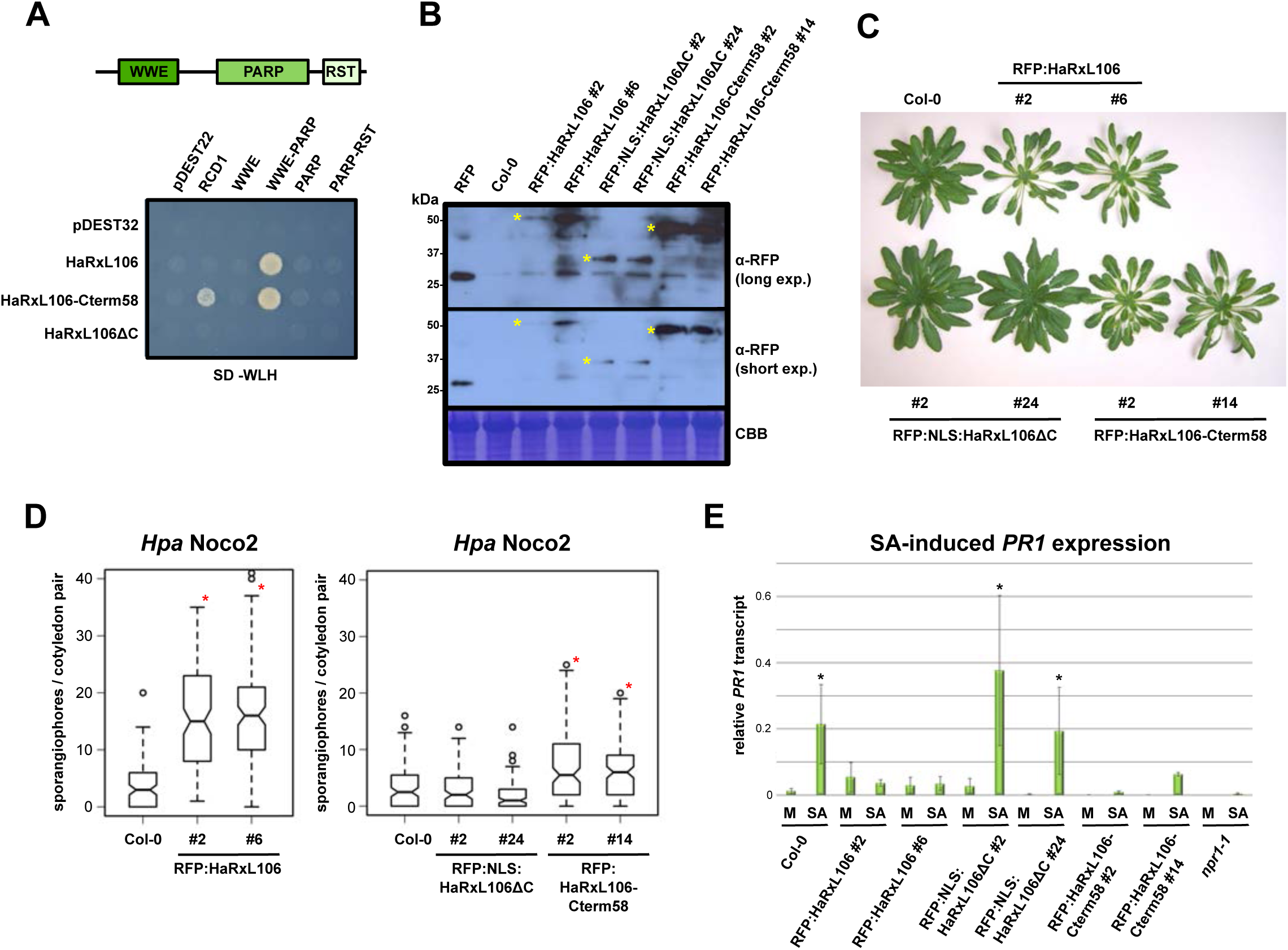
The C-terminal 58 amino acids of HaRxL106 are required for binding to RCD1 and mediate suppression of light and defense signaling. **(A)** Protein-protein interactions between RCD1, HaRxL106 and the indicated deletion constructs in a yeast-two-hybrid assay. pDEST32 and pDEST22 are empty bait and prey vectors, respectively. HaRxL106ΔC is a deletion construct lacking HaRxL106’s 56 C-terminal amino acids. HaRxL106-Cterm58 is an N-terminal deletion construct consisting only of the 58 C-terminal amino acids. **(B)** Western blot showing protein levels of RFP-tagged HaRxL106, NLS:HaRxL106ΔC and HaRxL106-Cterm58 in selected stable transgenic *Arabidopsis* lines. Asterisks indicate the expected migration based on the molecular weight of the fusion proteins. **(C)** Morphology of transgenic *Arabidopsis* lines expressing RFP-tagged HaRxL106, HaRxL106ΔC or HaRxL106-Cterm58. **(D)** Basal resistance to the virulent *Hpa* isolate Noco2 in 10-day-old seedlings of Col-0 and transgenics expressing RFP-tagged HaRxL106, HaRxL106ΔC or HaRxL106-Cterm58. The plots show the number of sporangiophores per cotyledon pair. Data from three independent biological experiments were pooled, horizontal bars show median, asterisks indicate mean values different from Col-0 (one-way ANOVA; Tukey-Kramer post-hoc test, p<0.05). **(E)** SA-induced *PR1* gene expression in Col-0, and transgenics expressing RFP-tagged HaRxL106, NLS:HaRxL106ΔC or HaRxL106-Cterm58. Four-week-old plants were sprayed with 0.1 mM SA or a mock solution and *PR1* expression levels were analyzed by qRT-PCR 8 h later. *PR1* expression levels were normalized by *EF1α* expression. The plot shows the mean of *PR1/EF1α* expression from three independent biological experiments. Error bars show SE, asterisks indicate mean values different from Col-0 mock treatment (one-way ANOVA; Tukey-Kramer post-hoc test, p<0.05).

As expected, transgenic lines expressing RFP-tagged HaRxL106 developed signs of constitutive shade avoidance (Fig. 6C). In contrast, RFP:NLS:HaRxL106 ΔC lines were indistinguishable from wild type plants. Notably, RFP-HaRxL106-Cterm58 lines resembled transgenics expressing full-length HaRxL106 suggesting that the C-terminal 58 amino acids of the effector are required and sufficient for attenuation of light signaling (Fig. 6C). We then tested resistance to *Hpa* Noco2 in these lines. While expression of RFP:HaRxL106 led to enhanced disease susceptibility in two independent transgenic lines (one-way ANOVA, F_2,342_=102.26, p=1.55 × 10^-35^; Tukey-Kramer post-hoc test, p<0.05), the RFP:NLS:HaRxL106 Δ C fusion protein failed to suppress defense. In contrast, transgenic lines expressing RFP:HaRxL106-Cterm58 were more susceptible to *Hpa* Noco2 than Col-0 (one-way ANOVA, F_4,590_=578.15, p=2.28 × 10^-25^; Tukey-Kramer post-hoc test, p<0.05), but less so than lines expressing the full-length effector. Therefore, the C-terminal 58 amino acids of HaRxL106 are required to alter light and defense signaling. Similar to RFP:HaRxL106 lines, transgenics expressing the HaRxL106 C-terminus responded with lower *PR1* transcript levels than wild type plants to SA spraying (Fig. 6E). In contrast, *PR1* transcript in lines expressing RFP:NLS:HaRxL106 ΔC responded like wild type plants to SA spraying (one-way ANOVA, F_15,32_=27.54, p=4.42 x 10^-14^; Tukey-Kramer post-hoc test, p<0.05).

### RCD1 is dispensable for basal resistance to *Hpa* but required for HaRxL106-mediated suppression of defense

To test if *RCD1* contributes to basal resistance to *Hpa,* we infected the *rcd1-1* mutant with *Hpa* Noco2. Despite the observed lower transcript abundance of NPR1 target genes in *rcd1-1* (Fig. 4), the mutant was not more susceptible than wild type plants (Fig. 7A). Consistent with a previous large-scale *Hpa* phenotyping report (Weßling et al., 2014), *rcd1-1* showed enhanced disease resistance compared to Col-0 (one-way ANOVA, F_5,1149_=79.22, p=1.70 × 10^-71^; Tukey-Kramer post-hoc test, p<0.05). This suggests that, if RCD1 is a relevant virulence target of HaRxL106, inhibition of RCD1’s function(s) or signaling is unlikely to be responsible for the enhanced disease susceptibility induced by HaRxL106. We considered that HaRxL106 may manipulate RCD1 in a way that is not mimicked by complete loss of RCD1 function, for example by altering RCD1’s interaction with other proteins or ligands. We compared susceptibility to *Hpa* Noco2 in the YFP:HaRxL106 line and a transgenic line expressing the same construct in an *rcd1-1* mutant background. Although the YFP:HaRxL106 fusion protein accumulated to similar levels in both transgenic lines (Fig. 7B), the *rcd1-1* mutation completely suppressed the enhanced disease susceptibility induced by YFP:HaRxL106 (Fig. 7A). To confirm that this effect is due to the *rcd1-1* mutation we re-introduced functional *RCD1* into the YFP-HaRxL106 *rcd1-1* background by crossing it with Col-0. Two independent F2 lines that were homozygous for *RCD1* and expressed the *YFP:HaRxL106* transgene were more susceptible to *Hpa* Noco2 than the parental YFP:HaRxL106 *rcd1-1* line (Fig. 7A; one-way ANOVA, F_5,1149_=79.22, p=1.70 x 10^-71^; Tukey-Kramer post-hoc test, p<0.05). Therefore, functional RCD1 protein is essential for suppression of basal defense by HaRxL106. Loss of *RCD1* function also attenuated the extent of petiole elongation in the YFP-HaRxL106 background (Fig. 7C; line #5) However, this effect was only partial with YFP:HaRxL106 *rcd1-1* #5 transgenics exhibiting an intermediate phenotype between *rcd1-1* and YFP:HaRxL106 in Col-0 background, perhaps because *SRO1* partially compensates for loss of *RCD1* function in such lines (Jaspers et al., 2009), or because HaRxL106 might target other host proteins in addition to RCD1.

**Figure 7.**
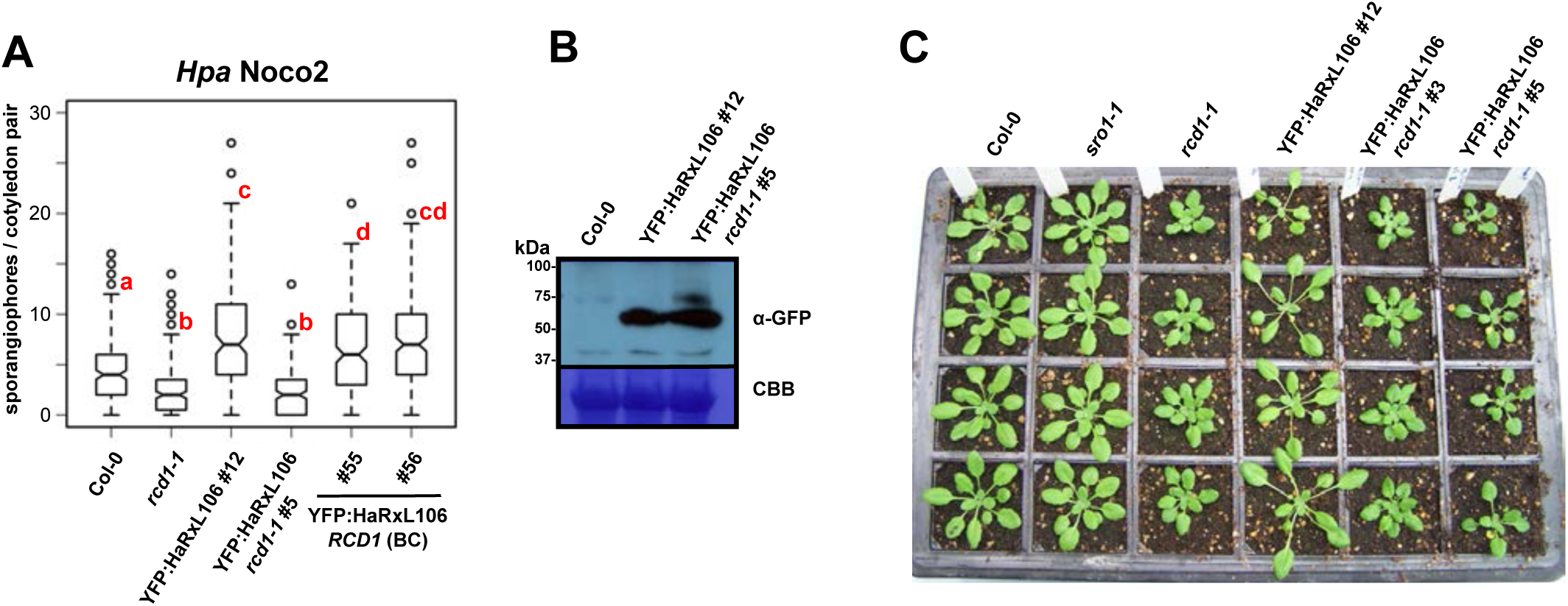
RCD1 is required for HaRxL106-induced suppression of basal defense. **(A)** Basal resistance to the virulent *Hpa* isolate Noco2 in 10-day-old seedlings of Col-0, *rcd1-1*, the YFP:HaRxL106 line #12, a transgenic lines expressing YFP:HaRxL106 to comparable levels in the *rcd1-1* background (#5) and two descendant lines of #5 in which RCD1 has been re-introduced by backcrossing to Col-0 (#55 and #56). The plots show the number of sporangiophores per cotyledon pair. Data from five independent biological experiments were pooled, horizontal bars show median, letters indicate differences between mean values (one-way ANOVA; Tukey-Kramer post-hoc test, p<0.05). **(B)** Western blot showing protein levels of YFP:HaRxL106 in line #12 (Col-0) and line #5 (*rcd1-1*). CBB = Coomassie brilliant blue stain. **(C)** Morphology of 4-week-old plants of Col-0, *sro1-1*, *rcd1-1*, and lines expressing YFP:HaRxL106 in Col-0 and *rcd1-1* backgrounds, respectively. The YFP:HaRxL106 fusion protein was not detectable by Western blot in line YFP:HaRxL106 *rcd1-1* #3.

### A crystal structure of RCD1’s PARP domain suggests that RCD1-type proteins do not function as canonical ADP-ribosyl transferases

Our findings that suppression of basal defense by HaRxL106 is largely dependent on RCD1 and that the effector binds to RCD1’s WWE-PARP domains prompted us to further investigate the molecular function(s) of RCD1. We reasoned that if RCD1 had PARP or a related transferase activity, HaRxL106 might manipulate this enzymatic function. To generate a structural framework for testing putative enzymatic functions of RCD1’s PARP domain we solved its structure by x-ray crystallography at 2.5 Å resolution (for data collection and refinement statistics see Table S2). As shown in Fig. 8A the RCD1 PARP domain adopts a fold that is overall similar to mammalian PARP domains. The closest structural homologue of the RCD1 PARP domain in the protein data bank was human PARP14 [PDB identifier 3SE2, in complex with inhibitor 6(5H)-phenanthridinone] with a root mean square deviation of 1.45 Å over 176 amino acids at a sequence identity of 17%. However, our structure of the RCD1 PARP domain also confirmed that the amino acid triad H-Y-E constituting the active site of mammalian and plant PARPs is not conserved in RCD1 (Kleine et al., 2008; Jaspers et al., 2010a) (Fig. 8B). The conserved Histidine (H1600) of HsPARP14 corresponds to RCD1 L333 whilst the position of Y1633 in HsPARP14 is filled by H365 of RCD1. These two first residues of the H-Y-E triad make direct contact to NAD^+^ and are essential for ADP-ribosyl transferase activity in mammalian enzymes (Kleine et al., 2008). The Glu residue of the H-Y-E triad that is critical for elongation of ADP-ribose chains, and therefore distinguishes mono-ADP-ribosyl-transferases from PARPs, is not conserved in HsPARP14. Consequently the enzymatic activity of HsPARP14 is limited to mono-ADP-ribosylation (Kleine et al., 2008). In the RCD1 PARP domain residue N428 takes the equivalent position. Non-conservation of the His and Tyr residues critical for NAD^+^ binding in canonical ADP-ribosyl-transferases suggests that the RCD1 PARP domain does not bind NAD^+^. Furthermore, the RCD1 PARP domain lacks an acidic residue in the third position of the triad. Overall, non-conservation of all of the residues critical for NAD^+^ binding and PARP activity strongly indicate that RCD1 does not have canonical PARP activity.

Many inhibitors of mammalian PARPs mimic the nicotinamide moiety of NAD^+^ and stabilize canonical PARP domains in thermal shift assays (Wahlberg et al., 2012). The PARP inhibitor 6(5H)-phenanthridinone binds to several human PARPs with affinity in the low micro-molar range (Wahlberg et al., 2012) and has also been proposed to bind to the corresponding cleft of RCD1 based on homology modelling (Rissel et al., 2017). To test whether 6(5H)-phenanthridinone also stabilizes RCD1 we compared thermal shifts induced by the inhibitor for the PARP domains of HsPARP1 and RCD1 (Fig. 8C). As previously shown, 6(5H)-phenanthridinone stabilized the catalytic domain of HsPARP1 (Wahlberg et al., 2012). In contrast the RCD1 PARP domain did not show any changes in stability in thermal shift assays, even in presence of 2 mM 6(5H)-phenanthridinone (Fig. 8C). This suggests that 6(5H)-phenanthridinone either does not bind to RCD1 or that the inhibitor binds in a way that is different from binding to human PARPs and therefore does not stabilize the RCD1 PARP domain.

Conceivably, the cleft of RCD1 that corresponds to the catalytic center of active PARPs has evolved to bind other small compounds. To unequivocally test if the cleft corresponding to the catalytic center of active PARPs is required for RCD1 function we designed structure-based site-directed mutants of RCD1 (Fig 8B). We mutated RCD1 residue H365 to Gln to abolish possible ring-stacking of small aromatic compounds, which is an important function of the corresponding Tyr (H-**Y**-E) residue in canonical PARPs. Furthermore, we mutated RCD1 S375 to a Trp. This mutation will abolish possible hydrogen bonding of the Ser residue with Nicotinamide-related compounds. In addition, we hypothesized that inserting a bulky hydrophobic residue at this position will occupy the cleft and block binding of potential ligands. Finally, we mutated RCD1 D421 to Ala. D421 hydrogen bonds to N428 and is the only acidic residue in close proximity that might possibly substitute for the lack of a Glu residue in RCD1 (Fig. 8B).

We transformed the RCD1 site-directed mutant constructs with a C-terminal triple HA epitope tag alongside with an RCD1-3HA wild-type control into the *rcd1-1* background. All constructs were under transcriptional control of 2.5 kb of the native *RCD1* promoter. As shown in Fig. 8D all constructs complemented the developmental phenotype of *rcd1-1*, demonstrating that the cleft of the RCD1 PARP domain, which corresponds to the catalytic site of active PARPs, is not essential for RCD1’s function in plant development. For one transgenic line expressing RCD1 D421A we found that while most plants showed wild type morphology, some plants were intermediate between wild type and *rcd1-1* (compare line B and B* in Fig. 8D). However, an independent transgenic line (A) expressing the same construct fully complemented *rcd1-1,* suggesting that differences in expression levels might account for partial complementation in line B. As a quantitative read-out for RCD1 function under stress conditions we determined the tolerance to oxidative stress induced by 1 µM Paraquat (Fig. 8E). The *rcd1-1* mutant was more resistant to Paraquat treatment than wild type plants, as previously reported (Ahlfors et al., 2004). The enhanced Paraquat tolerance of *rcd1-1* was reverted by expression of *RCD1*_*Pro*_*:RCD1:3HA*. Likewise, all mutant versions of RCD1 complemented the Paraquat tolerance phenotype of *rcd1-1* and we did not detect strong quantitative differences between *RCD1* and its mutant variants (Fig. 8E). For the RCD1 D412A construct line A fully complemented the enhanced Paraquat tolerance of *rcd1-1*. In contrast, line B that only partially complemented the developmental phenotype of *rcd1-1* did not complementation the Paraquat tolerance. Nevertheless, for each RCD1 mutant variant we identified at least one transgenic line that fully complemented both, *rcd1-1*‘s growth phenotype and the elevated tolerance to Paraquat. This demonstrates that mutations in the putative NAD^+^ binding site of RCD1 do not affect RCD1’s function in plant development and oxidative stress signaling. Taken together our results strongly suggest that RCD1 has no canonical PARP activity, does not bind to a broad-spectrum PARP inhibitor in thermal shift assays, and that the integrity of the presumed NAD^+^ binding cleft is not essential for RCD1’s functions in plant development and signal transduction under oxidative stress conditions.

### HaRxL106 binds to the N-terminal domains of RCD1 and SRO1 that mediate homo- and hetero-dimerization

Although RCD1 does not carry a canonical PARP domain, HaRxL106 appears to manipulate RCD1 function by binding to the WWE-PARP region of RCD1 (Fig. 6A). Using the Y2H system we further narrowed down the HaRxL106 binding site of RCD1 to an N-terminal fragment encompassing the WWE domain and the linker region up to the beginning of the PARP domain (Fig. 9A). Deletion of the linker region resulted in loss of interaction with HaRxL106 suggesting that the WWE domain on its own is not sufficient for binding to the effector. The isolated PARP domain did not bind to HaRxL106, irrespective of whether or not we included the linker region (Fig. 9A). HaRxL106 showed binding to a construct containing WWE, linker and PARP domain but to a lower extent than binding to the WWE-linker region (Fig. 9A). In contrast, the HaRxL106ΔC control did not interact with any of the RCD1 deletion constructs tested. This suggests that the WWE-linker region is required and sufficient for binding to HaRxL106.

WWE domains are thought to mediate protein-protein interactions (Aravind, 2001). We therefore tested if the WWE-linker regions of RCD1 and SRO1 can mediate formation of homo- or hetero-oligomers. We found that the RCD1 WWE-linker region interacts with itself and the corresponding region of SRO1 in Y2H assays indicative of the formation of homo- and hetero-oligomers. We obtained comparable results for the corresponding part of the SRO1 protein (Fig. 9C). These data suggest that the RCD1 and SRO1 WWE-linker regions could mediate formation of RCD1/SRO1 oligomers.

### RCD1’s WWE domain forms protein complexes with histone modifying kinases

Given that RCD1 does not have PARP activity, we further characterized RCD1 protein function(s) by screening for *in planta* interactors of RCD1. Attempts to immuno-purify epitope-tagged RCD1 protein in amounts sufficient for LC-MS/MS analysis of co-purifying proteins from transient expression assays in *N. benthamiana* or stable *Arabidopsis* transgenics were not successful. We therefore resorted to screening for interactors of RCD1’s WWE-linker region following transient expression in *N. benthamiana* as this part of the protein binds to HaRxL106 and is more stable (Fig. 9D). The predominant interactors were several importin-α isoforms, full-length RCD1-type proteins and protein kinases with sequence homology to Casein II kinases (Fig. 9E; Table S3). RCD1 and related proteins carry conserved NLS motifs at their N-termini (Katiyar-Agarwal et al., 2006) and co-purification of importin-α proteins confirms that these NLS bind to nuclear transport adapters in plant cells. Given that we identified several peptides from the PARP and RST domains (Fig. S2) of RCD1-type proteins by LC-MS/MS, the WWE-linker fragment appears to form homo- and hetero-oligomers with endogenous RCD1-type proteins in *N. benthamiana*, which is consistent with oligomer formation of the WWE-linker regions in Y2H (Fig. 9B and 9C). Our pull-down strategy therefore provides an indirect method to immuno-purify full-length RCD1-type proteins. Another prominent group of proteins co-precipitating with the WWE-linker fragment were Casein II-like kinases. A blastp search against the *Arabidopsis* protein database (TAIR11) identified MUT9-like kinases (MLKs) as likely orthologs (Fig. 9E). To our knowledge MLKs have not been characterized in *N. benthamiana.* MLKs are nuclear-localized er/Thr inases hat hosphorylate istones. n *Arabidopsis* and*Chlamydomonas* phosphorylation of histone H3 Thr3 (H3T3ph) is the best characterized MLK phosphorylation site (Casas-Mollano et al., 2008; Wang et al., 2015). We found several residues in the RCD1 WWE linker region to be phosphorylated (Fig. S3; Table S4). While most of these phosphorylated peptides were located in the GFP:WWE-linker bait protein from *Arabidopsis,* we also detected two phospho-peptides from the WWE-PARP linker region of a co-purifying *N. benthamiana* RCD1 ortholog (Fig. S3), indicating that RCD1-type proteins might be MLK substrates. Overall our results show that MLKs interact with RCD1-type proteins in plant cells, suggesting a possible role of RCD1 and sequence-related proteins in influencing covalent modifications of histone tails.

To test protein-protein interactions in *Arabidopsis* we also purified the WWE-linker domain from a stable transgenic *Arabidopsis* line expressing *35S*_*Pro*_*:GFP:WWE-linker* protein. Protein levels immuno-purified from *Arabidopsis* were substantially lower compared to those obtained in *N. benthamiana* experiments. Nevertheless, in a single experiment we identified unique peptides of the four *Arabidopsis* MLKs in immuno-precipitates of the WWE-linker region and confirmed that the WWE-linker region is phosphorylated (Fig. S3; Table S5). In accordance with the results from *N. benthamiana* experiments we also identified several peptides from the PARP and RST domains of RCD1 and SRO1 (Fig. S2).

MLKs have been previously reported to affect H3T3ph levels in response to osmotic and salt stress and the *mlk1,2* double mutant is hypersensitive to sub-lethal concentrations of PEG and NaCl (Wang et al., 2015). As MLKs and RCD1 form protein complexes in plant cells, and given that SA marker genes are expressed at lower levels in *rcd1* mutants, we asked if MLKs also affect the transcriptional response to SA. We sprayed *mlk1,2,3* and *mlk1,3,4* triple mutants with SA and determined transcriptional upregulation of *PR1* 8 h later (Fig. 9F). The *mlk1,3,4* triple mutant consistently showed elevated *PR1* transcript levels in response to SA (one-way ANOVA, F_3,8_=79.22, p=0.01; Tukey-Kramer post-hoc test, p<0.05), whereas basal *PR1* expression levels in the mock control were not strongly altered. In contrast the *mlk1,2,3* triple mutant responded like wild type, suggesting that MLK4 might be particularly relevant for SA-induced transcript changes.

## Discussion

Several biotrophic pathogens evolved virulence mechanisms to counteract activation of SA-dependent defense genes (Asai et al., 2014; Lewis et al., 2015), but how pathogen effectors suppress SA-mediated immunity remains only partially understood. Apart from active conversion of SA (Djamei et al., 2011), effector-mediated activation of JA signaling appears to be the main strategy of biotrophic pathogens to attenuate SA-dependent defense (Zheng et al., 2012; Caillaud et al., 2013; Yang et al., 2017). Here we show that *Hpa* effector HaRxL106 induces shade avoidance in *Arabidopsis* and suppresses SA signaling while JA marker genes are either not affected or slightly repressed (Table S1).

We mapped both, the light signaling- and defense-manipulating activities of HaRxL106 to a short C-terminal part of the effector. Intriguingly, loss of *RCD1* function renders *Arabidopsis* resilient to HaRxL106-mediated suppression of defense and diminishes HaRxL106-induced petiole elongation (Fig. 7). These results suggest that RCD1 integrates signals downstream of both, pathogen- and photoreceptors and that *Hpa* is exploiting this function of RCD1 to attenuate plant immunity. Notably, the morphological and defense phenotypes of *rcd1* null mutants are opposite to those induced by ectopic expression of HaRxL106 (Fig. 7). Therefore, it is conceivable that the effector manipulates RCD1 to boost its cellular activities although this hypothesis cannot easily be tested without a better understanding of the molecular function of both proteins. HaRxL106 is predicted to have an α -helical WY structure, a fold which likely evolved as a versatile building module of oomycete effectors and can mediate different molecular functions in fusion with small peptides or other domains (Boutemy et al., 2011; Maqbool et al., 2016). The WY domain in HaRxL106 might function as a scaffold that stabilizes and/or presents the C-terminal peptide that is essential for suppression of plant immunity. In accordance with this model, expressing a fusion of the C-terminal 58 amino acids of HaRxL106 to RFP is sufficient to trigger shade avoidance and attenuate basal defense (Fig. 6). Notably, manipulation of selective autophagy by the host-targeted *Phytophthora infestans* effector PexRD54 is also based on a disordered C-terminal peptide that is stabilized by five tandem WY domains (Dagdas et al., 2016; Maqbool et al., 2016).

RCD1, and sequence-related proteins from *Arabidopsis* and rice, bind transcription factors via their C-terminal RST domains. In contrast the function(s) of RCD1’s N-terminal WWE and central PARP domains have not been characterized. Although an RCD ortholog from wheat shows PARP activity (Liu et al., 2014), our structural analysis suggests that *Arabidopsis* RCD1 is unlikely to be enzymatically active. Our crystal structure of the RCD1 PARP domain provides first insights into plant PARP domains and we identified several molecular differences between RCD1’s PARP domain and the catalytic domain of mammalian PARPs. None of the amino acids constituting the H-Y-E triad that are essential for PARP activity in previously characterized enzymes are conserved in RCD1 (Fig. 8B). Our complementation analysis of RCD1 site-directed mutant variants shows that RCD1 tolerates relatively severe amino acid exchanges in the cleft region of its PARP domain without losing its functions in plant development and oxidative stress signaling (Fig. 8D and E). Taken together, our structural and molecular analyses of RCD1 suggest that the protein does not have PARP activity and can therefore, in analogy to pseudo-kinases, be classified as a pseudo-PARP. Given that the N- and C-terminal domains of RCD1 mediate protein-protein interactions, the PARP domain of RCD1 might act as a scaffold bridging and/or coordinating the action of the terminal protein interaction domains.

**Figure 8.**
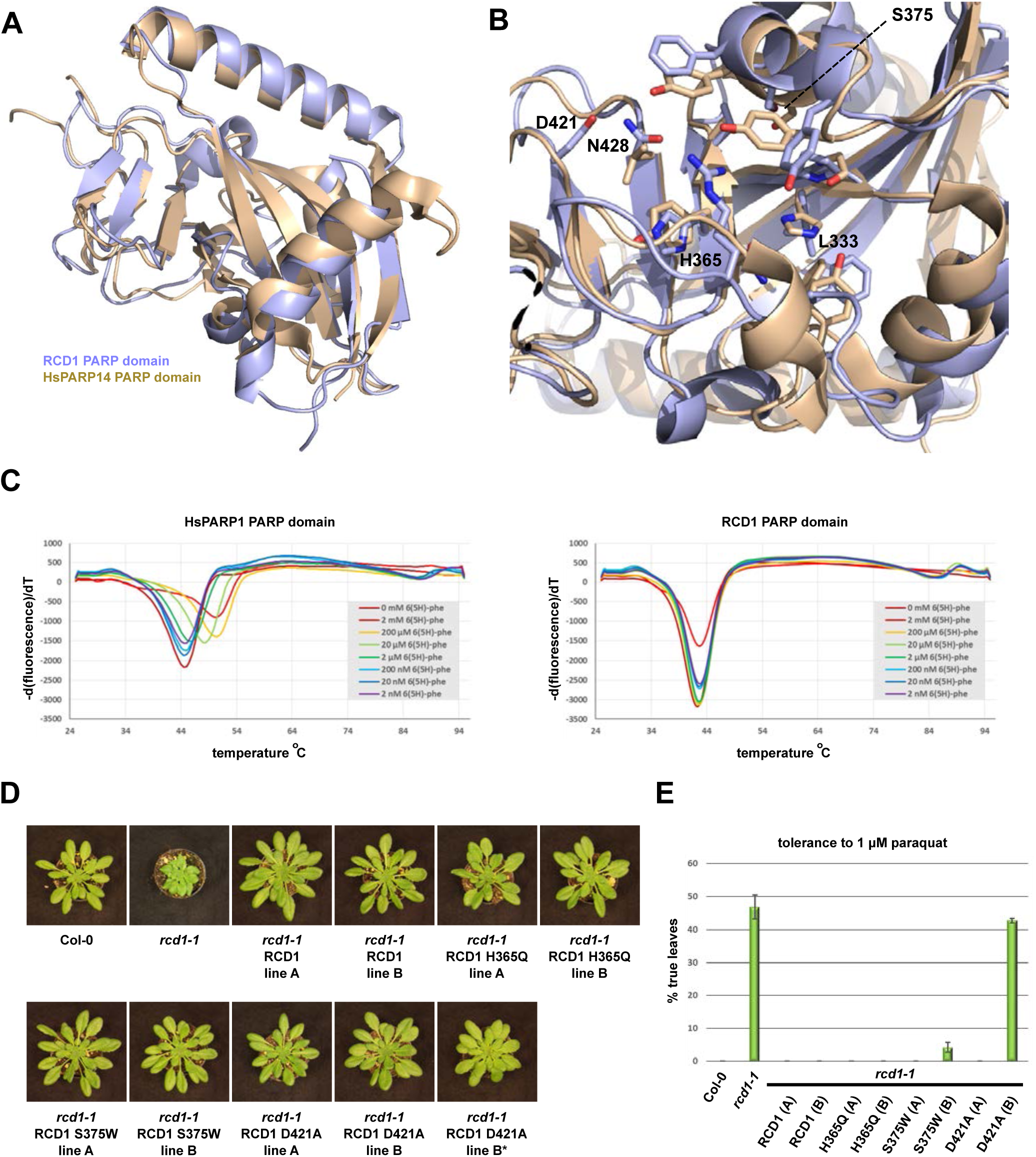
Mutations in RCD1’s putative NAD^+^ binding pocket do not compromise RCD1 function in development and oxidative stress signaling. **(A)** Crystal structure of the RCD1 PARP domain (light blue, PDB code 5NGO) superimposed onto the structure of HsPARP14 (beige, PDB code 3SE2; Wahlberg et al., 2012). **(B)** Structural comparison of HsPARP14’s NAD^+^ binding site (beige) with the corresponding area of the RCD1 PARP domain (light blue). RCD1 residues L333, H365 and N428 take the positions of the conserved H-Y-E triad in canonical PARPs. **(C)** Effect of the non-specific PARP inhibitor 6(5H)-phenanthridinone on the thermal stability of HsPARP1’s PARP domain (left panel) and the RCD1 PARP domain (right panel). **(D)** Morphological phenotypes of RCD1 site-directed mutants with the indicated amino acid exchanges in the putative NAD^+^ binding site. Images are representative of >10 plants per line. RCD1 D421A line B showed both, plants that fully complemented and individuals that partially complemented (asterisk) the developmental phenotype of *rcd1-1*. **(E)** Tolerance of Col-0, *rcd1-1* and RCD1 site-directed mutants to Paraquat. Seeds were sown on MS plates containing 1 μM Paraquat and the percentage of seedlings (n=48) that had developed true leaves after 20 days was determined. The plot shows the mean value of two independent biological experiments. Error bars show SE.

RCD1’s WWE domain and the linker region up to the PARP domain are essential for binding to HaRxL106 (Fig. 9A). We also found that the WWE domains of RCD1 and its paralog SRO1 can form homo- and hetero-oligomers in Y2H (Fig. 9). It will be interesting to clarify if oligomer formation has a biological function in signal transduction of RCD1-type proteins. The WWE domain is a conserved iso-ADP-ribose binding domain but it is not known if plant WWE domains bind PAR chains (He et al., 2012; Wang et al., 2012). Conceivably, HaRxL106 binding to RCD1’s WWE domain could interfere with ADP-ribose binding if this biological function is conserved in plants. An alternative but not mutually exclusive scenario is that RCD1’s WWE domain is an interaction module for other proteins. Here, we identified kinases from the MLK group as novel interactors of RCD1’s WWE-linker domain. Consistent with complex formation between the N-terminal domain of RCD1 and MLKs we identified several phosphorylation sites in RCD1’s linker region (Supplementary Table S3). The interaction between MLKs and the RCD1 N-terminus implies that phosphorylation of the linker region might be mediated by MLKs, but our data do not rule out alternative kinases. The ∼90 amino acid linker region between the RCD1 WWE and PARP domains is predicted to be disordered (Ishida and Kinoshita, 2007; Kragelund et al., 2012) but it is conceivable that phosphorylation or binding of interacting proteins induce a specific fold in this region (Wright and Dyson, 2009; Bah et al., 2015). As the RCD1 PARP domain appears to act as a scaffold, reversible phosphorylation of residues in the linker region between WWE and PARP domains could regulate the cooperation of these two domains.

**Figure 9.**
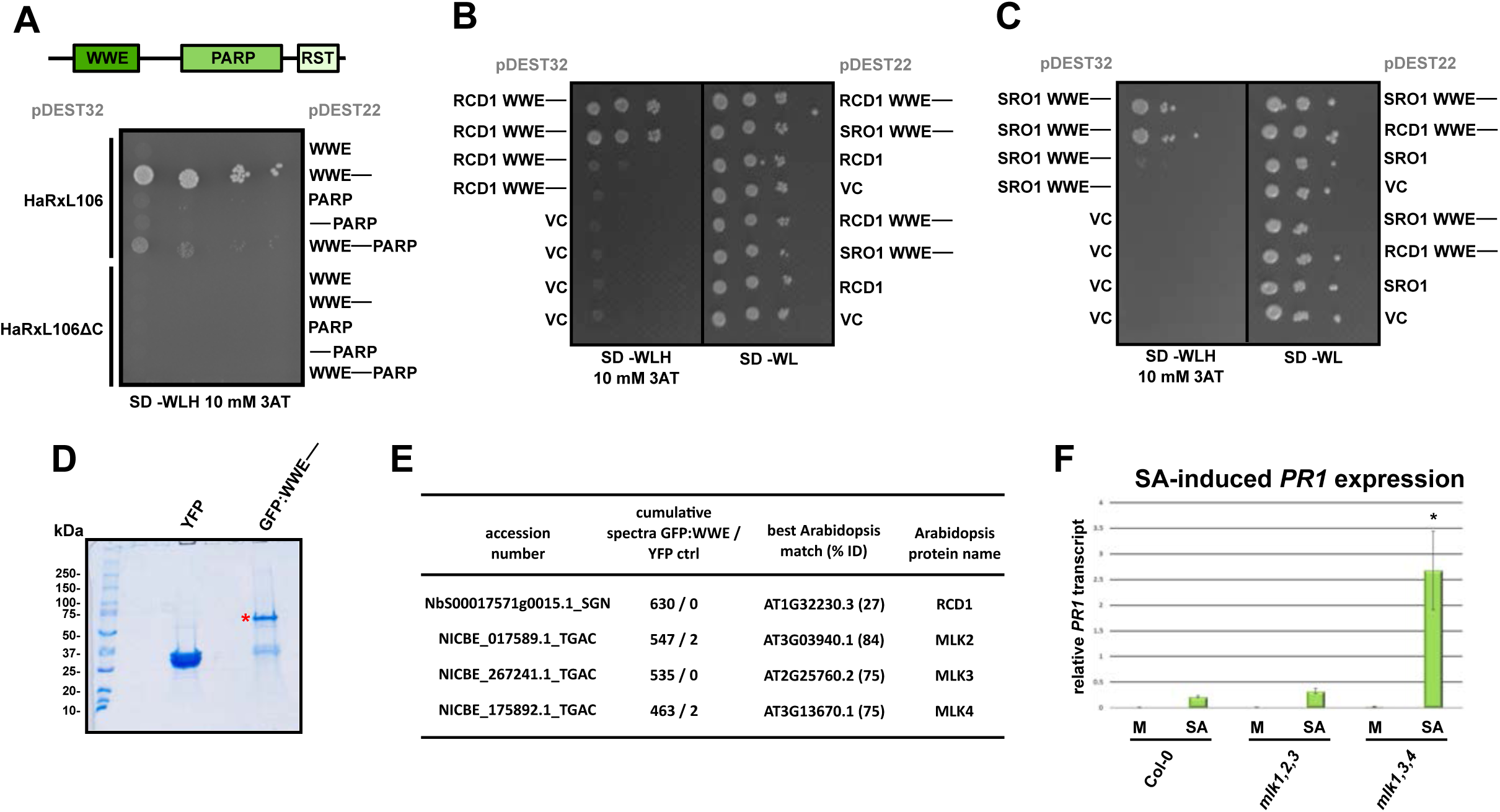
The N-terminal WWE-linker region of RCD1 forms homo- and hetero-oligomers and interacts with Mut9-like kinases (MLKs). **(A)** Mapping protein-protein interactions between HaRxL106 and the indicated RCD1 domain(s) in the yeast-two-hybrid assay. The HaRxL106ΔC deletion construct does not bind RCD1 and is shown as a control. **(B)** and **(C)** Formation of homo- and hetero-oligomers by RCD1’s (B) and SRO1’s (C) WWE-linker region in the yeast-two-hybrid assay. VC = pDEST22 or pDEST32 vector control. **(D)** Representative SDS-PAGE image of transiently expressed YFP and RCD1 GFP:WWE-linker proteins immuno-purified from *N. benthamiana*. The gel was stained with colloidal blue Coomassie. The asterisk indicates the expected migration of the GFP:WWE-linker fusion protein. **(E)** Selected proteins that were enriched in immunoprecipitates of the GFP:WWE-linker fusion protein compared to the YFP control. For the full list see Supplementary Table S3. **(F)** SA-induced *PR1* gene expression in Col-0 and the *mlk1,2,3* and *mlk1,3,4* triple mutants. Four-week-old plants were sprayed with 0.1 mM SA or a mock solution and *PR1* expression levels were analyzed by qRT-PCR 8 h later. *PR1* expression levels were normalized by *EF1α* expression. The plot shows the mean of *PR1/EF1α* expression from three independent biological experiments. Error bars show SE, asterisk indicates mean value different from Col-0 SA treatment (one-way ANOVA; Tukey-Kramer post-hoc test, p<0.05).

MLKs are recruited to the evening complex in a PHYB-dependent manner and have previously reported functions in light signaling, circadian rhythm and abiotic stress responses (Casas-Mollano et al., 2008; Wang et al., 2015; Huang et al., 2016). Here we show that an *mlk1,3,4* triple mutant responds with higher expression of the SA marker gene *PR1* to exogenous application of SA while basal *PR1* levels are unaffected (Fig. 9F). This suggests that MLK phosphorylation sites on histones or other proteins also influence transcriptional mechanisms required for fine-tuning the amplitude of SA-induced *PR1* expression. H3T3 is the best-characterized phosphorylation site of MLKs in *Chlamydomonas* and *Arabidopsis*. Notably, in mammalian cells repressive H3T3ph and activating tri-methylation of the adjacent K4 form a molecular switch that directly affects TFIID binding thereby regulating gene expression in different phases of the cell cycle (Varier et al., 2010). In mammals the TAF3 subunit anchors the TFIID complex onto H3K4me3-marked nucleosomes (Vermeulen et al., 2007; van Ingen et al., 2008). In contrast, the TAF3 ortholog of plants, if it exists, has not been identified (Lawit et al., 2007; Fromm and Avramova, 2014). Consistent with H3T3ph being a repressive chromatin mark, we find that simultaneously knocking out *MLK1, 3, and 4* results in higher SA-induced *PR1* expression.

Although attenuated defense responses under shade conditions have been reported for multiple plant-pathogen or plant-herbivore interactions, the underlying molecular mechanisms are only partially understood (Ballaré, 2014). Notably, constitutive SA defense mutants still undergo shade avoidance, arguing against simple resource partitioning scenarios to explain the ‘growth-defense tradeoff’ (de Wit et al., 2013). Reciprocally, the *sav3-2* mutant that is impaired in the morphological responses to shade still shows attenuated activation of JA-dependent pathogen resistance under low R/FR but it remains unknown if the same holds true for SA-dependent immunity (Cerrudo et al., 2012). Of note, unlike HaRxL106 over-expressors, plants infected by (hemi-)biotrophic pathogens do not show morphological signs of shade avoidance. We hypothesize that this discrepancy is explained by the tissue-unspecific expression of the *HaRxL106* transgene by the *35S* promoter. In contrast, *Hpa* primarily infects epidermal and mesophyll leaf cells in natural infections (Koch and Slusarenko, 1990).

Overall, these observations indicate that activation of at least one signaling pathway under shade, rather than the morphological responses triggered by it, are responsible for suppression of innate immunity. PIF transcription factors integrate light, temperature and other environmental signals (Wigge, 2013). Recently, PIF4 was shown to act as a negative regulator of plant immunity at elevated temperature (Gangappa et al., 2016). Constitutive defense activation and the dwarf stature of the Nod-like receptor mutant *snc1-1* is suppressed when plants are grown at elevated temperature, and this depends on PIF4. A similar effect was observed for overexpression of *PHYB* in the *snc1-1* mutant background (Gangappa et al., 2016). Interestingly, also mutations in *RCD1* render *snc1-1* insensitive to elevated temperature although this effect was only observed for plant size and not for disease resistance to the bacterial strain *Pst* DC3000 (Zhu et al., 2013). Nevertheless, the similar phenotypes observed in *snc1-1 pif4*, *snc1-1 35S:PHYB* and *snc1-1 rcd1* double mutants indicate that RCD1 might be a component of a regulatory node integrating light, temperature and defense signals. RCD1 interacts with PIF transcription factors and *rcd1* mutants show reduced hypocotyl elongation under red and blue light (Jaspers et al., 2009; Salazar, Felipe Sarmiento and Neuhaus, Gunther, 2010). Although the molecular function(s) of RCD1 remain poorly characterized, its localization to the nucleus and interaction with transcription factors point to a role as a transcriptional co-regulator. Consistent with such a role of RCD1 is our finding that HaRxL106 interferes with SA signaling at the level of transcription (Fig. 3).

In summary, our analysis of an *Arabidopsis* downy mildew effector, HaRxL106, has helped us reveal previously unsuspected roles of the RCD1 family of proteins in plant immunity, and a likely contribution of the MLK family of protein kinases to defense signaling.

## Methods

### Plants and growth conditions

For hypocotyl growth assays *Arabidopsis* seeds were surface sterilized, sown on plant growth medium [Mourashige Skook medium incl. MES and vitamins (Duchefa # M0255), 0.1 g/l Myoinositol, 8 g/l Bactoagar], stratified for 48 h at 4 °C in the dark and subsequently germination was induced by a 6 h white light stimulus. The plates were placed in long-day (12 h light / 12 h dark) conditions at 21 °C and the fluence rate of white light was adjusted to 12 µmol m^-2^ s^-1^ by placing layers of Whatman filter paper above the plates. We determined hypocotyl length on day 5 using by photographing flattened seedlings and measuring hypocotyl length using ImageJ software 1.43u (Wayne Rasband, National Institutes of Health). Growth conditions for *N. benthamiana* and all other *Arabidopsis* experiments were as in Fabro et al. (2011) and Segonzac et al. (2011). The *mos6-1*, *asil1-1*, *rcd1-1*, *rcd1-3* mutants have been described (Palma et al., 2005; Gao et al., 2009; Jaspers et al., 2009). The *mlk1,2,3* and *mlk1,3,4* triple mutants have been described in Huang et al. (2016).

### Generation of transgenic *Arabidopsis* lines

Transgenic *Arabidopsis* plants expressing YFP–tagged HaRxL106 have been described in Wirthmueller et al. (2015). To generate transgenic lines expressing HS:HaRxL106, RFP:HaRxL106, RFP:NLS:HaRxL106ΔC and RFP:HaRxL106-Cterm58 (all from the *Hpa* Emoy2 allele of HaRxL106) we used the following previously described pENTR plasmids: pENTR4-HaRxL106, M followed by HaRxL106 amino acids I^25^-S^285^ (Fabro et al., 2011), pENTR/D-TOPO-SV40NLS:HaRxL106ΔC, sequence APKKKRKV followed by HaRxL106 amino acids I^25^-G^229^, and pENTR/D-TOPO-HaRxL106-Cterm58, HaRxL106 amino acids G^228^-S^285^ (Wirthmueller et al., 2015). Plasmids pXCSG-HS and pH7WGR2 were recombined with the above pENTR plasmids using Gateway^®^ LR clonase II to generate HS- and RFP-tagged versions of the HaRxL106 constructs, respectively. Transgenic HS-tagged HaRxL106 lines were generated by transforming Col-0 or *35S*_*Pro*_*:NPR1:GFP* plants with *A. tumefaciens* strain GV3101 pMP90^RK^ carrying pXCSG-*HS:HaRxL106* constructs. Transgenic lines expressing RFP-tagged HaRxL106 constructs were generated by transforming Col-0 with *A. tumefaciens* strain GV3101 pMP90 carrying the corresponding pH7WGR2 plasmids. We transformed plants using floral dip (Logemann et al., 2006). Col-0 lines expressing RCD1 amino acids 1-265 as C-terminal fusion to GFP were generated by recombining a corresponding pENTR clone with pK7WGF2 (Karimi et al., 2002), transforming the construct into *A. tumefaciens* strain GV3101 pMP90 and subsequently transforming *Arabidopsis* Col-0 plants by floral dip. Site-directed mutants of RCD1 were generated using the QuikChange method (Agilent). To generate site-directed mutants of RCD1 a 428 bp BamHI/NcoI fragment was subcloned from the RCD1 coding sequence into the BamHI/NcoI sites of pET-DUET. Following successful mutagenesis using the QuikChange method, BamHI/NcoI fragments were cloned into a pENTR/D-TOPO plasmid carrying 2499 bp of the *RCD1* promoter sequence fused to the *RCD1* coding sequence. The pENTR/D-TOPO plasmids carrying wild-type or mutated RCD1 were then recombined with pGWB13 (Nakagawa et al., 2007) to generate a translational fusion with a triple HA-tag at the C-terminus of the protein. The constructs were transformed into the *rcd1-1* mutant by floral dip as described above.

### *Hpa* infection and quantification

Basal resistance to *Hpa* isolate Noco2 was either tested on adult leaves of 6-week-old plants (Fig. 1A) or on the cotyledons of 10-day-old seedlings grown on soil. For both types of experiments plants were sprayed with a suspension of 1 × 10^5^ *Hpa* Noco2 spores per ml. The plants were placed in high (>90%) humidity under a plastic dome. Sporulation on seedlings was scored at 5 days post infection, sporulation of adult plants was quantified 7-8 days post infection. For the adult leaf assay 20 leaves per genotype were harvested and stained with Trypan blue. Following destaining with chloral hydrate solution conidiophores on 20 leaf areas of 1 cm^2^ were counted using a light microscope. For the seedling assay 35-40 seedlings per genotype were incubated in a 0.02% (w/v) solution of Uvitex 2B, then destained in water for 2 min, mounted on a Styrofoam rack and imaged through a Leica UV filter (Leica #10447415) using a Leica M165 FC fluorescent stereo microscope connected to a EL6000 laser source. Only conidiophores on the upper side of the cotyledons were counted.

### Quantification of SA

Six-weeks-old *Arabidopsis* plants were syringe-infiltrated with 1 × 10^8^ cfu/ml *Pst* DC3000 in 10 mM MgCl_2_ or a 10 mM MgCl_2_ mock solution. Samples were taken 24 h after infiltration including control samples from non-treated plants. SA was extracted and quantified as previously described (Aboul-Soud et al., 2004). Briefly, leaf tissues (0.2 g) were extracted in 1 ml 90% methanol following homogenization in liquid nitrogen. 3-hydroxybenzoic acid (HBA, Sigma) was used as an internal standard. The level of SA was determined by a fluorescence detector (Shimadzu RF-20AXS, excitation at 305 nm and emission at 405 nm) in reverse-phase HPLC using a Nexera UHPLC (Shimadzu) system with a C18 (Kinetex 2.6µm XB-C18) column.

### SA treatment

For SA treatment *Arabidopsis* plants were grown on soil under short day conditions. One h after dawn (09:00) we sprayed 4-week-old plants with a solution containing 0.1 mM SA and 0.01% Silwet L-77 until run-off and took samples for RNA preparations 8 h later.

### Paraquat treatment

*Arabidopsis* seedlings were sown on GM medium supplemented with 1 µM Paraquat and placed in long-day conditions. After 14 days the number of seedlings with expanded true leaves was counted.

### RNA extraction, cDNA synthesis and RNA-Seq transcriptome profiling

Five-week-old *Arabidopsis* plants were syringe-infiltrated with 5 × 10^5^ cfu/ml of *P. syringae* DC3000 at 12:00 (4 h after lights on). Rosette leaf samples from non-treated, mock- and bacteria-infiltrated plants were harvested 24 h later. Total RNA was extracted using the TRI reagent (Sigma) and 1-Bromo-3-chloropropane (Sigma), as per manufacturer’s guidelines. RNA was precipitated with half volume of isopropanol and half volume of high salt precipitation buffer (0.8 M sodium citrate and 1.2 M sodium chloride). RNA samples were treated with DNaseI (Roche) according to the manufacturer’s recommendation and phenol/chloroform extracted and ethanol precipitated. For qRT-PCR assays, mRNA was reverse transcribed using SuperScript II Reverse Transcriptase (Thermo Fisher) according the manufacturer’s conditions. cDNA samples were diluted fivefold and 1 µl was used as template in a 20 µl qRT-PCR reaction. For RNA-seq, 3 µg of total RNA was used to generate first strand cDNA using a oligo(dT) primer comprising the P7 sequence of Illumina flow-cell. Double strand cDNA was synthesized as described previously (Okayama and Berg, 1982). Purified cDNA was subjected to Covaris shearing to a target size of 200bp (parameters: Intensity – 5, Duty cycle – 20%, Cycles/Burst – 200, Duration – 90seconds). End repairing and A-tailing of sheared cDNA was carried out as described by Illumina. Y-shaped adapters were ligated to A-tailed DNA and subjected to size selection on 1x TAE agarose gels. The gel-extracted library was PCR enriched and quantified using qPCR with previously sequenced similar size range Illumina library. Transcriptome data has been deposited at NCBI’s Gene Expression Omnibus (<https://www.ncbi.nlm.nih.>gov/geo/) with identifier GSE89402.

### Yeast-two-Hybrid

All constructs for the Y2H assay were cloned into pDEST32 (bait) or pDEST22 (prey) vectors (Invitrogen) using Gateway® recombination. The RCD1, WWE-PARP (amino acids 1-471) and PARP-RST (amino acids 241-589) deletion constructs used for Fig. 6A have been published (Jaspers et al., 2009). Two additional constructs, WWE (amino acids 1-155) and PARP (amino acids 247-472), were cloned into pDEST22 using the same strategy. For the Y2H assays in Fig. 9 A-C the following RCD1 and SRO1 deletion constructs were generated: RCD1 WWE (amino acids 1-170), RCD1 WWE--- (amino acids 1-265), RCD1 PARP (amino acids 265-460), RCD1 ---PARP (amino acids 170-460), RCD1 WWE---PARP (amino acids 1-460), SRO1 WWE--- (amino acids 1-262). For HaRxL106 constructs the pENTR plasmids carrying HaRxL106, HaRxL106ΔC or HaRxL106-Cterm58 were recombined into pDEST32. Yeast strain Mav203 was co-transformed with pDEST32 and pDEST22 plasmids and double-transformed yeast cells were selected on SD –Leu –Trp plates. Selected clones were grown in liquid SD –Leu – Trp medium at 30 °C for 48 h until cultures had reached saturation. The OD_600_ was adjusted to 0.1, and the yeast strains were plated in 10-fold serial dilutions onto SD –Leu –Trp –His medium containing 0, 1, 5, 10 or 20 mM 3-AT. Serial dilutions were also plated on SD –Leu –Trp medium to compare growth rate of the yeasts under non-selective conditions. The plates were photographed 3-5 days after plating.

### qRT-PCR

qRT-PCR was performed using 10 µl SYBR Green qPCR premix (Sigma) and the following oligonucleotides: EF1α-fw CAGGCTGATTGTGCTGTTCTTA, EF1α-rv GTTGTATCCGACCTTCTTCAGG, PR1-fw ATGAATTTTACTGGCTATTCTC, PR1-rv AGGGAAGAACAAGAGCAACTA. qRT-PCR reactions were run in duplicates or in triplicates on a CFX96 Touch− Real-Time PCR Detection System (Bio-Rad) using the following program: (1) 95 °C, 3 min; (2) [95 °C for 30 s, 60 °C for 30 s, 72 °C for 30 s] ×41, (3) 72 °C for 10 min followed by (4) a melting curve analysis from 55 °C to 95 °C. The data were analyzed using CFX Manager− software (Bio-Rad). The average relative transcript levels from three independent biological replicates were analyzed using one-way ANOVA followed by a Kramer-Tukey post hoc test.

### Transient expression

RCD1:GFP and SRO1:GFP constructs were generated by cloning their coding sequences into pENTR/D-TOPO and recombining these plasmids with pK7FWG2 (Karimi et al., 2002). *A. tumefaciens* GV3101 strains were grown on selective plates, resuspended in 10 mM MgCl_2_ 10 mM MES pH 5.6 and incubated with 100 µM acetosyringone for 2 h at RT. Prior to infiltration each strain was mixed with *A. tumefaciens* strain GV3101 expressing the silencing suppressor 19K at a ratio of 1:2[19K]. For co-expression, strains were mixed in a 1:1:2[19K] ratio. We infiltrated leaves of 4-week-old *N. benthamiana* plants with a needleless syringe and harvested the leaves 48–72 h later.

### Protein extraction from *Arabidopsis*, *N. benthamiana*, immunoprecipitation and western blot

Protein extracts were prepared by grinding *N. benthamiana* or *Arabidopsis* leaf material in liquid nitrogen to a fine powder followed by resuspension in extraction buffer [50 mM Tris, 150 mM NaCl, 10% glycerol, 1 mM EDTA, 5 mM DTT, 1× protease inhibitor cocktail (Sigma #P9599), 0.2% NP-40, pH 7.5] at a ratio of 2 ml buffer per 1 g leaf material. For experiments that included RCD1 (domains) the proteasome inhibitor MG132 (10 µM) and phosphatase inhibitors NaF (10 mM) and Na_3_VO_4_ (1 mM) were added to all buffers. Crude protein extracts were centrifuged at 20.000 x g 4°C 20 min and the supernatant was either boiled in sodium dodecyl sulphate (SDS) sample buffer for western blots or used for immunoprecipitation. For western blots proteins were separated by SDS-PAGE and electro-blotted onto polyvinylidene difluoride membrane (Millipore). Antibodies α-HA 3F10 (Roche), α-GFP 210-PS-1GFP (Amsbio), α-RFP-biotin ab34771 (Abcam) were used for detection. For immunoprecipitation a fraction of the supernatant was saved as ‘input’ sample and 15 µl GFP-beads (GFP-Trap_A; Chromotek) were added to the remaining supernatant. The volume for immunoprecipitation from *N. benthamiana* for western blots was 1.4 ml. The volume for immunoprecipitation from *Arabidopsis* was 4 ml. The volume for immunoprecipitation experiments coupled to mass spectrometry was 15-25 ml. Following incubation of the samples on a rotating wheel at 4°C for 2 h the beads were collected by centrifugation at 1200 x g and 4°C for 1.5 min. The beads were washed 3 times with 1 ml extraction buffer and then boiled in SDS sample buffer to elute proteins from the beads.

### Thermal shift assays

Thermal shift assays were performed in a CFX96 Touch− Real-Time PCR Detection System (Bio-Rad) as described by Vivoli et al. (2014) with minor modifications. Briefly, 23 µl reactions containing the PARP domains of HsPARP1 [L713F mutant, Langelier et al. (2012)] or RCD1 at a final concentration of 0.11 mg/ml were prepared in reaction buffer [40 mM HEPES pH7.5, 150 mM NaCl, 6.6x SYPRO Orange (Thermo Fisher)]. 6(5H)-phenanthridinone (Sigma) was added from a 25 mM stock solution in DMSO to final concentrations of 2 nM to 2 mM or DMSO was used as a control. The total reaction volume was 25 µl. Unfolding of the proteins was determined by running a thermal denaturation program with 0.5 °C temperature increase per cycle followed by measuring SYPRO orange fluorescence in FRET mode of the instrument. The first derivative of the fluorescence over temperature was plotted using CFX Manager− software (Bio-Rad).

### Confocal microscopy

*Arabidopsis* leaf discs were mounted onto microscopy slides in 50% glycerol or water and analyzed on a Leica DM6000B/TCS SP5 confocal microscope with the 488 nm as excitation wavelengths for GFP.

### Recombinant expression and purification of PARP domains from *E. coli*

The expression construct for the human PARP1 PARP domain (amino acids 662-1014; L713F mutant) has been described (Langelier et al., 2012). The PARP domain of RCD1 (amino acids 269-460) was cloned into the pOPINF vector (Berrow et al., 2007) linearized by KpnI/HindIII digest using Gibson assembly. The expression construct was transformed into SoluBL21 DE3 *E. coli* cells (Genlantis, NEB). For protein expression four 1 l cultures were grown in LB medium at a temperature of 37 °C to an OD_600_ of 1.0 – 1.2. The cultures were cooled to 18 °C before expression was induced by the addition of 0.5 mM IPTG for 16 h. Cells were pelleted by centrifugation (5000 x g, 4 C, 12 min) and the pellets were resuspended in buffer A (50 mM Tris-HCl, 0.3 M NaCl, 20 mM imidazole, 5% glycerol, 50 mM glycine, pH 8.0) supplemented with 0.1% Polyethylenimine and 1x cOmplete− EDTA-free protease inhibitor cocktail (Roche). Cells lysis was induced by addition of lysozyme (1 mg/ml final concentration, RT, 15 min) followed by sonication. Insoluble proteins and cell debris were removed by centrifugation (30.000 x g, 4 C, 20 min) and the supernatant was loaded onto a 5 ml HisTrap HP IMAC column (GE Healthcare). The column was washed with buffer A until the A_280_ reached 25 mAU and proteins were eluted using buffer B (50 mM Tris-HCl, 0.3 M NaCl, 20 mM imidazole, 5% glycerol, 50 mM glycine, 500 mM Imidazole, pH 8.0). The elution from the IMAC column was injected onto a size exclusion chromatography column [Superdex 75 26/60 PG column (GE Healthcare) pre-equilibrated in 20 mM HEPES, 150 mM NaCl, pH 7.5]. PARP domains eluting from the column were concentrated to 0.5-1 mg/ml (HsPARP1) and used for thermal shift assays. For crystallization of the RCD1 PARP domain, the His6-tag was cleaved using 3C protease. The protein was run through a 5 ml HisTrap HP IMAC column in buffer A1 to remove the His6 tag and residual uncleaved fusion protein, followed by injection onto the Superdex 75 26/60 PG column and eluted as above. The protein was concentrated to 35 mg/ml using Vivaspin 20 and 2 columns (Sartorius) with a molecular weight cut-off of 5 kDa and flash-frozen in liquid nitrogen. Seleno-Met-labelled RCD1 PARP protein was produced using feedback-inhibition and purified as described above for native RCD1 PARP.

### Crystallization and data collection

The RCD1 PARP domain was used for crystal screens at a concentration of 35 mg/ml. Crystals of native and Seleno-Met-labelled protein grew in 0.1 M HEPES pH7.5, 1.6 M Ammoniumsulfate, 2% PEG1000 in hanging drops at 20 °C. The crystals were harvested in Paratone-N oil (Hampton) and frozen in liquid nitrogen. Data collection was performed on beamlines i04 (native) and i24 (Seleno-Met) at Diamond Light Source, Oxford, UK. X-ray data were processed with iMosflm (Battye et al., 2011) and scaled with Aimless (Evans and Murshudov, 2013) from the CCP4 suite (Collaborative Computational Project, Number 4, 1994). For X-ray data collection statistics see Table S2. The RCD1 PARP structure was solved by single wavelength anomalous dispersion using Phaser (McCoy, 2007). Iterative building and refinement cycles with Coot (Emsley et al., 2010), Refmac5 (Murshudov et al., 2011) and Phenix (Adams et al., 2010) were used to obtain the final model with statistics given in Table S2. Validation tools in Molprobity (Chen et al., 2010) and Coot were used to analyze the final structure. 3D visualizations of protein structures were prepared using PyMOL software v1.7.2 (http://sourceforge.net/projects/pymol/). Reflection data and the RCD1 PARP domain structure have been deposited at the Protein Data Bank with identifier 5NGO.

### Protein mass spectrometry

Samples for LC-MS analysis were prepared by excising bands from one dimensional SDS-PAGE gels stained with colloid Comassie Brilliant Blue (Instant Blue, Expedeon). The gel slices were destained with 50% Acetonitrile and cysteine residues modified by 30 min reduction in 10 mM DTT followed by 20 min alkylation with 50 mM chloroacetamide. After extensive washing with 30% Acetonitrile and dehydration with 100% acetonintrile the pieces were incubated with 100 ng of trypsin (Promega) in 100 mM ammonium bicarbonate, 10% acetonitile at 37 °C overnight.

LC-MS/MS analysis was performed using a hybrid mass spectrometer Orbitrap Fusion and a nanoflow UHPLC system U3000 (Thermo Scientific). The generated peptides were applied to a reverse phase trap column (Acclaim Pepmap 100, 5 µm, 100 µm x 20 mm) connected to an analytical column (Acclaim Pepmap 100, 3µm, 75µm x 500mm; Thermo Scientific). Peptides were eluted in a gradient of 9-50% acetonitrile in 0.1% formic acid (solvent B) over 50 min followed by a gradient of 50-60% B over 3 min at a flow rate of 300 nL min^-1^. The mass spectrometer was operated in positive ion mode with nano-electrospray ion source with an ID 0.01 mm fussed silica PicoTip emitter (New Objective). Voltage +2.2 kV was applied via conductive T-shaped coupling union. Transfer capillary temperature was set to 320 °C, no sheath gas, and the focusing voltages were in factory default setting. Method MS events consisted from full scan in Orbitrap analyzer followed by two collisions of “softer” CID and more “energetic” HCD to maximize the chances to acquire spectra with structurally important information. Orbitrap full scan resolution of 120000, mass range 300 to 1800 m/z automatic gain control (AGC) target 200000 and maximal infusion time 50ms were set. Data-dependent algorithm MS/MS fragmentation of large number of precursor ions was used. The dynamic exclusion 30 s, “Top speed” precursor selection method within 3 s cycle between full scans and “Universal method” for infusion time and AGC calculation were used. The isolation width 1.6 m/z and normalized collision energy of 30% were set for both CID and HCD collisions. Only the precursor ions with positive charge states 2 – 7 and intensity greater than 10000 were selected for MS/MS fragmentation.

### Software processing and peptide identification

Peak lists in the form of Mascot generic files (mgf files) were prepared from raw data using MS Convert (Proteowizard project) and sent to peptide match search on the Mascot server using Mascot Daemon (Matrix Science). Peak lists were searched against *N. benthamiana* or *A. thaliana* databases. Tryptic peptides with up to 2 possible miscleavages and charge states +2, +3, +4 were allowed in the search. The following modifications were included in the search: oxidized methionine (variable), carbamidomethylated cysteine (static). Data were searched with a monoisotopic precursor and fragment ion mass tolerance 10 ppm and 0.6 Da respectively. Mascot results were combined in Scaffold (Proteome Software) and exported to Excel (Microsoft Office) for sample-to-sample comparison. In Scaffold, the peptide and protein identifications were accepted if probability of sequence match and protein inference exceeded 95.0% and 99%, respectively. At least 2 identified peptides per protein were required. Protein probabilities were calculated by the Protein Prophet algorithm; proteins that contained similar peptides and could not be differentiated based on MS/MS analysis alone were grouped to satisfy the principles of parsimony (Searle, 2010).

## Acknowledgements

We thank Marcel Wiermer (University of Göttingen) for critically reading the manuscript. We thank the BBSRC-UK (grants: BBJ00453, BBK009176), the Gatsby Charitable Foundation, the John Innes Foundation, Dahlem Centre of Plant Sciences and the German Research Foundation (DFG; grant WI 3670/2-1) for funding. JK and MW are members of the Centre of Excellence in the Molecular Biology of Primary Producers (2014-2019), which is funded by the Academy of Finland (decisions #271832 and #307335). Research in the MW laboratory is funded by the Academy of Finland (decisions #275632 and #283139). SA acknowledges funding from Japan Society for the Promotion of Science, grant KAKENHI15K1865/17K07679. We thank Jeff Dangl (University of North-Carolina, Chapel Hill), Jim Beynon (University of Warwick), Pascal Braun (Helmholtz Centre Munich) and Marc Vidal (Harvard) for sharing data on HaRxL106-interacting proteins. We thank Heidrun Häweker (TSL) for propagating *Hpa*, Torsten Schultz-Larsen (TSL) for help with establishing the *Hpa* Uvitex 2B assay on cotyledons, Ram Krishna Shrestha (TSL) for help with figure preparation in *R*, and Tuomas Puukko for performing the Y2H analysis in Fig. 6. We thank the Diamond Light Source (beamlines I04 and I24 under proposal MX7641) for access to their X-ray data collection facilities. We thank Lionel Hill (JIC Metabolomics department) for sharing access to HPLC equipment. LW would like to thank Tina Romeis (FU Berlin) for providing lab space and sharing equipment. This work was supported by a FEBS long-term fellowship awarded to LW (2010-2013) and a RIKEN Special Postdoctoral Fellowship awarded to SA.

## Author contributions

Conceived and designed the experiments: LW, SA, GR, GF, DK, MW, JK, MB, JJ

Performed the experiments: LW, SA, GR, JS, GF, DK, RL, MW

Analyzed the data: LW, SA, GR, JS, GF, DK, MW, DM, MB

Contributed reagents/materials/analysis tools: GR, GF, MW, JK, DM, FM, MB, JJ

Wrote the paper: LW

All authors have edited and approved the submitted version of the manuscript.

---

**Supplementary Figure S1.**
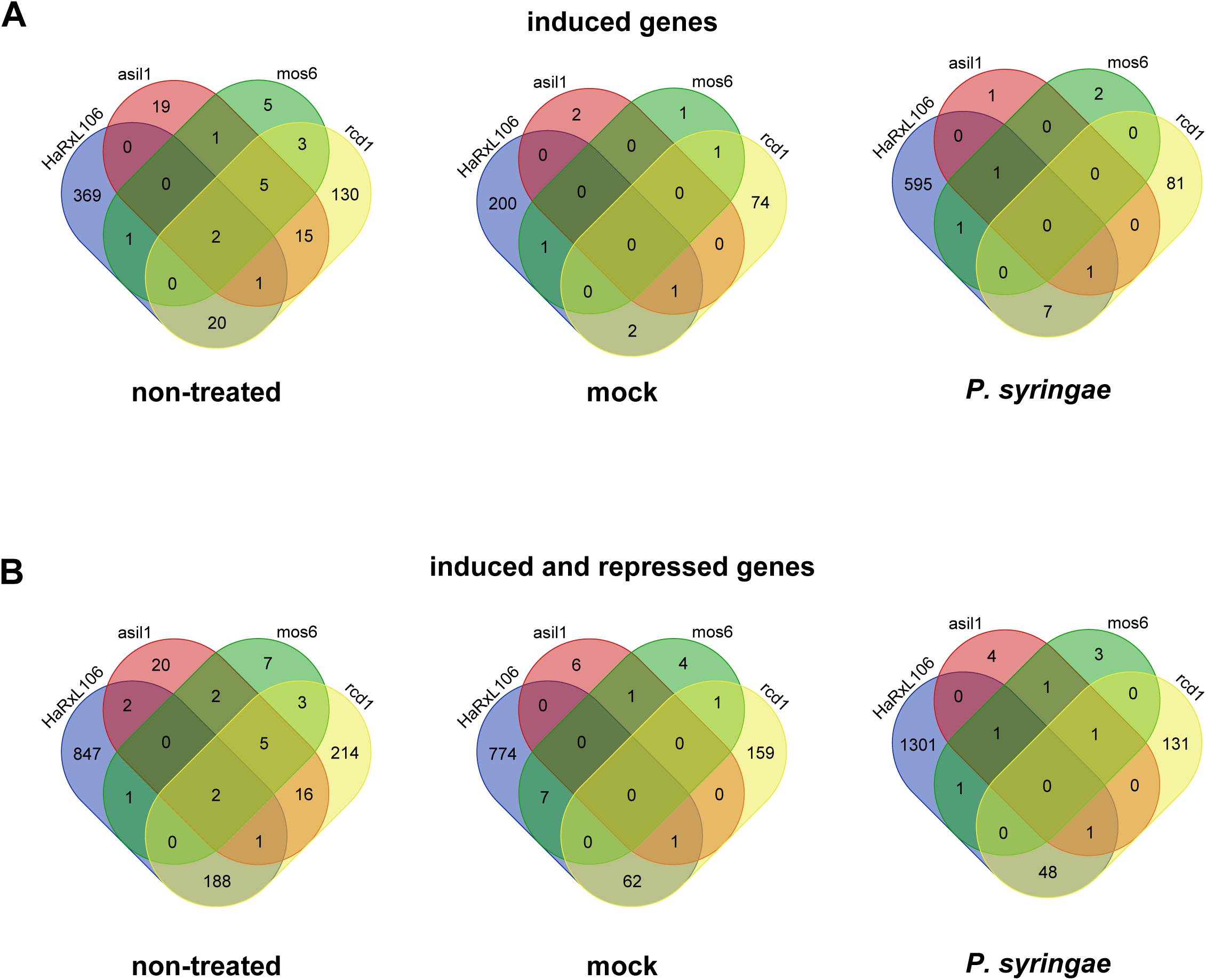
Overlap of genes differentially expressed between HS:HaRxL106 and Col-0 and the transcriptome profiles of *mos6*, *asil1* and *rcd1*. Upper panel shows genes that were over-expressed in the HS:HaRxL106 line #2 compared to Col-0. Lower panel shows all genes differentially expressed between HS::HaRxL106 line #2 and Col-0. Venn diagrams were prepared using http://bioinformatics.psb.ugent.be/webtools/Venn/.

**Supplementary Figure S2.**
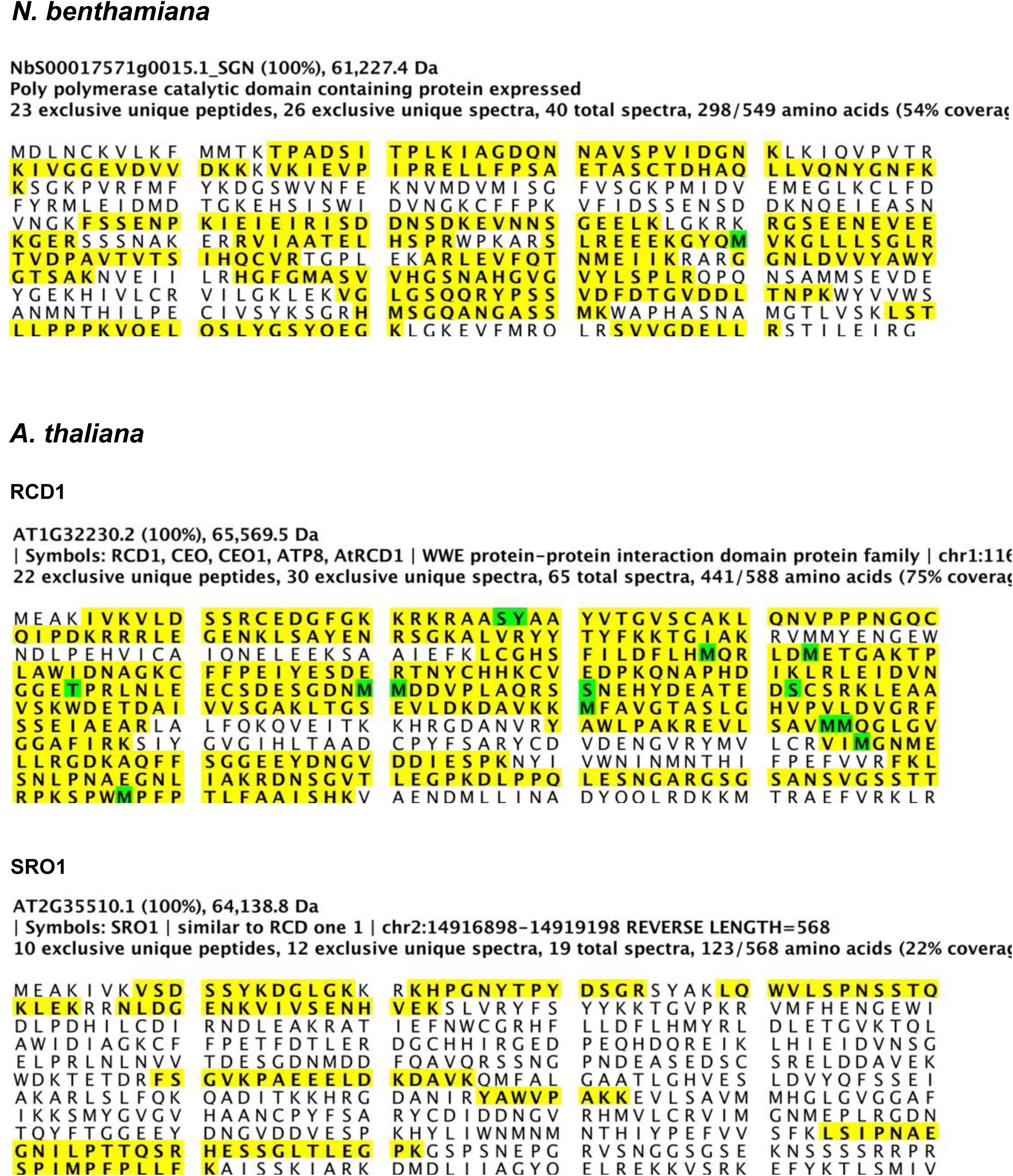
Sequence coverage of NbRCD1, AtRCD1 and AtSRO1 identified by peptide fingerprinting in immunoprecipitation experiments of the *Arabidopsis* RCD1 GFP:WWE-linker fusion protein. Tryptic peptides are shown in yellow. Green color indicates modified peptides (S/T/Y possible phosphorylation sites; M artificial oxidation or carbamidomethylation).

**Supplementary Figure S3.**
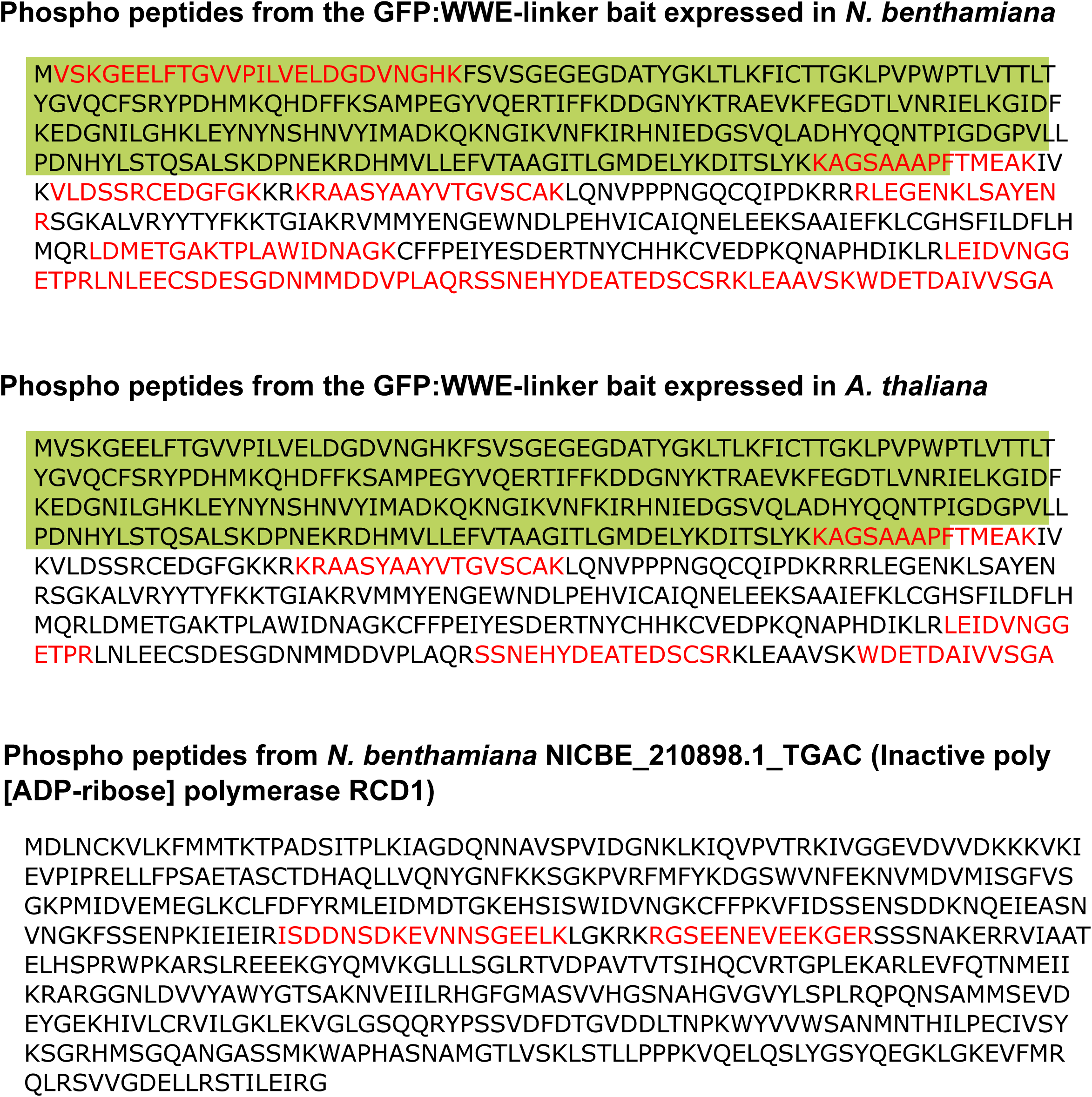
Phospho peptides identified in the *Arabidopsis* RCD1 GFP:WWE-linker fusion protein and a co-purifying NbRCD1 ortholog. The GFP sequence is indicated in green and phospho peptides are shown in red. For NbRCD1 three MS spectra supporting the two identified phosho peptides are shown. For a list of all identified phospho peptides from the *N. benthamiana* experiments including peptide identification probabilities and Mascot ion scores see Table S4.

**Supplementary Table S1.** Lists of DEGs in non-treated, mock- or *Pst*-infiltrated leaf tissue and analysis of GO terms and cis-regulatory elements in genes differentially expressed in *35S*_*Pro*_*:HS:HaRxL106* or *rcd1-1*.

**Supplementary Table S2.** X-ray data collection, refinement, and validation statistics.

**Supplementary Table S3.** List of proteins identified in immunoprecipitates of the RCD1 WWE-linker bait protein from *N. benthamiana*. Data are from three independent biological experiments, each consisting of two independently processed samples of GFP:WWE-linker and one or two YFP control samples.

**Supplementary Table S4.** Phospho peptides identified in immunoprecipitates of the RCD1 WWE-linker bait protein from *N. benthamiana*.

**Supplementary Table S5.** List of proteins identified in the immunoprecipitate of the RCD1 WWE-linker bait protein from *A. thaliana.*

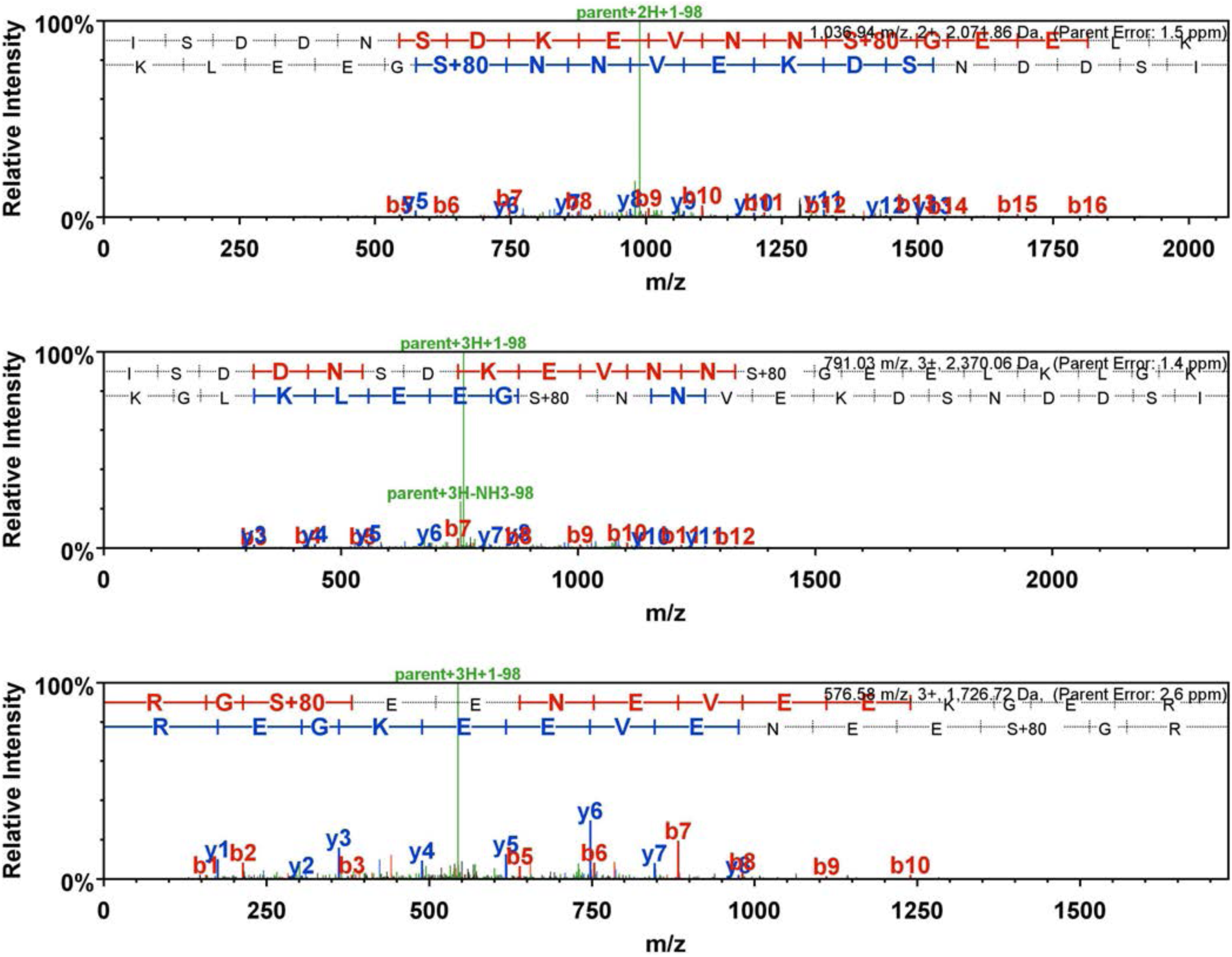
MS spectra of phospho peptides from N. benthamiana NICBE_210898.1_TGAC (Inactive poly [ADP-ribose] polymerase RCD1)

